# TRIBAL: Tree Inference of B cell Clonal Lineages

**DOI:** 10.1101/2023.11.27.568874

**Authors:** Leah L. Weber, Derek Reiman, Mrinmoy S. Roddur, Yuanyuan Qi, Mohammed El-Kebir, Aly A. Khan

## Abstract

B cells are a critical component of the adaptive immune system, responsible for producing antibodies that help protect the body from infections and foreign substances. Single cell RNA-sequencing (scRNA-seq) has allowed for both profiling of B cell receptor (BCR) sequences and gene expression. However, understanding the adaptive and evolutionary mechanisms of B cells in response to specific stimuli remains a significant challenge in the field of immunology. We introduce a new method, TRIBAL, which aims to infer the evolutionary history of clonally related B cells from scRNA-seq data. The key insight of TRIBAL is that inclusion of isotype data into the B cell lineage inference problem is valuable for reducing phylogenetic uncertainty that arises when only considering the receptor sequences. Consequently, the TRIBAL inferred B cell lineage trees jointly capture the somatic mutations introduced to the B cell receptor during affinity maturation and isotype transitions during class switch recombination. In addition, TRIBAL infers isotype transition probabilities that are valuable for gaining insight into the dynamics of class switching.

Via *in silico* experiments, we demonstrate that TRIBAL infers isotype transition probabilities with the ability to distinguish between direct versus sequential switching in a B cell population. This results in more accurate B cell lineage trees and corresponding ancestral sequence and class switch reconstruction compared to competing methods. Using real-world scRNA-seq datasets, we show that TRIBAL recapitulates expected biological trends in a model affinity maturation system. Furthermore, the B cell lineage trees inferred by TRIBAL were equally plausible for the BCR sequences as those inferred by competing methods but yielded lower entropic partitions for the isotypes of the sequenced B cell. Thus, our method holds the potential to further advance our understanding of vaccine responses, disease progression, and the identification of therapeutic antibodies.

**Availability:** TRIBAL is available at https://github.com/elkebir-group/tribal

## 1 Introduction

B cells play a pivotal role in the adaptive immune response, producing antibodies that neutralize foreign substances and infections [1, 2]. These antibodies, consisting of a heavy chain and a light chain, are initially formed as sequence-specific B cell receptors (BCRs) (Fig. 1a). The generation of specific BCR genes stems from a process known as V(D)J recombination, where DNA segments are rearranged to ensure a wide spectrum of antibodies to counter various pathogens [3]. To enhance their effectiveness, B cells undergo affinity maturation (Fig. 1b) [4], a a micro-evolutionary process involving repeated cycles of *somatic hypermutation* (SHM) and cellular divisions. SHM introduces mutations in the BCR genes, selecting for B cells expressing high-affinity BCRs, while eliminating those with low affinity. Con-currently, B cells have the ability for *class switch recombination* (CSR) (Fig.1b) [5], which diversifies their response by altering the antibody’s functional class or *isotype*.

**Figure 1.**
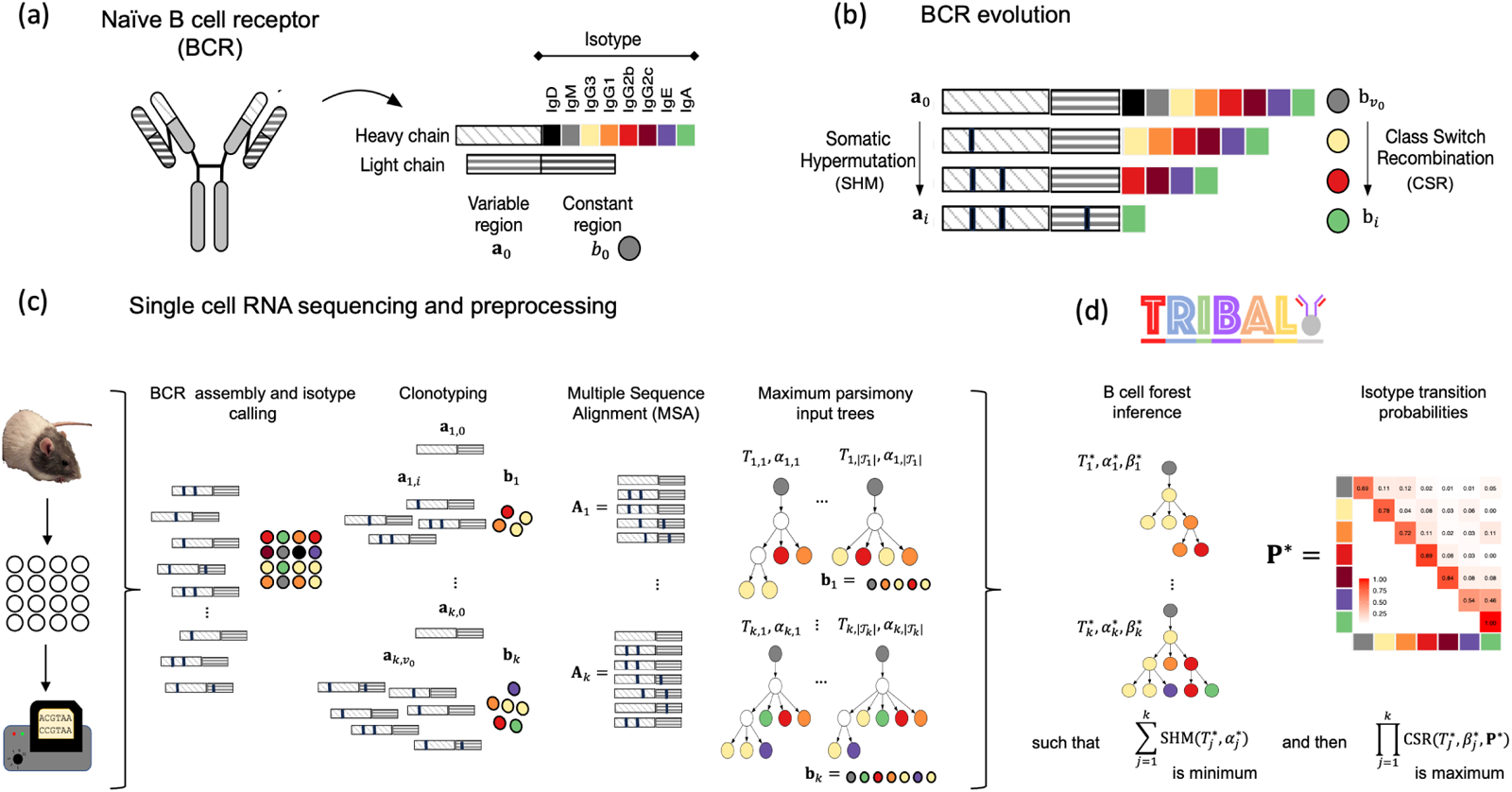
TRIBAL infers B cell lineage trees and isotype transition probabilities for scRNA-seq data. (a) A BCR consists of paired heavy and light immunoglobulin chains, each consisting of a variable and constant region. The isotype is the heavy chain constant locus that is transcribed. (b) The BCR of B cells undergo somatic hypermutation/affinity maturation, where point mutations are introduced into the variable region of the heavy and light chains, and class switch recombination, where the heavy chain constant locus undergoes recombination and beings transcribing a different isotype. (c) After scRNA-seq, the variable regions for the light and heavy immunoglobulin alleles are assembled, the isotypes are called and the B cells are clustered into *k* clonotypes. A multiple sequence alignment **A***j* is found for each clonotype *j* and used to infer a set of input trees with maximum parsimony. The leafs of each input tree are labeled by isotypes **b**. (d) TRIBAL jointly infers a B cell lineage tree 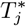 for each clonotype *j* and population-specific isotype transition probabilities **P**^*\**^ with maximum parsimony for MSA **A***j* and maximum likelihood for isotypes **b***j*.

Understanding the evolutionary history of B cells during these adaptive processes is integral to better understanding B cell response to infection and vaccination. However, the selection pressures applied to B cells during the affinity maturation process necessitate more specialized analytical approaches than those utilized for species phylogeny inference [6–9]. Specifically, Hoehn et al. developed HLP17 [10] and HLP19 [11], which are specialized codon substitution models for use with maximum likelihood inference via IgPhyML. Another important difference between B cell and species evolution is the lower mutation rate and the relatively short length of the BCR sequence (*≈* 600 bp). These properties imply that maximum parsimony inference methods are viable [12–14] but typically result in a large solution space of many plausible phylogenies for the same data. In addition, these solutions exhibit additional tree uncertainty in the form of *polytomies*, i.e., multifurcating nodes with more than two children. This is counter to the underlying cell lineage tree, which is bifurcating as cell division results in exactly two daughter cells.

One approach to resolve phylogenetic uncertainty is the inclusion of additional data. This approach has proven effective in other areas, such as the use of physical location for studying cancer migration and metastasis [15], or geographical location for the inference of gene flow [16]. More related to B cell lineage inference, sequence abundance has been utilized by both GCTree [12] and ClonalTree [17] to discriminate between candidate solutions. However, both methods were originally developed for bulk RNA sequencing data and consequently were restricted to analysis of only the heavy chain sequences. With single cell RNA-sequencing (scRNA-seq), it is now possible to efficiently assemble BCR sequences that includes both heavy and light chain from a population of B cells [18] (Fig. 1c). As a result, the evolution of BCRs during affinity maturation can now be tracked with scRNA-seq with higher fidelity [19]. In addition, scRNA-seq yields another valuable data source in the form of the expressed isotype of the constant region of the heavy chain (Fig. 1c) [19]. Isotype data extracted from scRNA-seq has been helpful in related immunological domains, such as inferring dynamic cellular trajectories [20]. However, isotype expression is an especially useful marker of B cell evolution. When a B cell undergoes class switching from its current isotype to a new isotype, any heavy chain constant region locus between the current isotype and the new isotype in the genome is cut out or removed via a recombination process (Fig. 1b). Consequently, CSR is an irreversible process and the isotype state of a B cell offers a distinct milestone in its evolutionary history. Therefore, the inclusion of isotype information into the problem of B cell lineage inference has the potential to help minimize phylogenetic uncertainty by both reducing the size of the solution space and yielding more refined B cell lineage trees with fewer polytomies.

In this work, we present TRIBAL, which stands for Tree Inference of B cell Clonal Lineages (Fig. 1d). TRIBAL utilizes both the BCR sequence and isotype information from sequenced cells to infer a B cell lineage tree that jointly models the evolutionary processes of SHM and CSR. Additionally, TRIBAL infers the underlying isotype transition probabilities providing valuable insight into the dynamics of CSR (Fig. 1d). We demonstrate the accuracy of TRIBAL on simulated data and show that it is effective on experimental single cell data generated from the 5’ 10x Genomics platform. TRIBAL is available open source and has the potential to improve understanding of vaccine responses, track disease progression, and identify therapeutic antibodies.

## 2 Problem Statement

Suppose that we have sequenced and subsequently aligned the variable regions of the heavy and light chain of the B cell receptor (BCR) of *n* B cells that descend from the same naive B cell post V(D)J recombination, resulting in a multiple sequence alignment (MSA) **A** composed of *m* columns — the typical alignment length is *m≈* 650 (Fig. 1c). That is, for each B cell *i*, we are given the concatenated sequence **a**_*i*_ *∈* Σ^*m*^ where Σ ={ *A, G, C, T*,} −. In addition, we are given the isotype *b*_*i*_ *∈* [*r*] = {1, …, *r*}, determined using tools such as Cell Ranger [21]. For humans, there are *r* = 8 isotypes ordered as IgM/D, IgG3, IgG1, IgA1, IgG2, IgG4, IgE and IgA2, whereas for mice there are *r* = 7 isotypes ordered as IgM/D, IgG3, IgG1, IgG2b, IgG2c (2a), IgE, IgGA. Finally, we are given sequence **a**_0_ and isotype *b*_0_ = 1 of the ancestral naive B cell of the *n* B cells — note that **a**_0_ and *b*_0_ is post V(D)J recombination but prior to somatic hypermutation (SHM) and class switch recombination (CSR) and thus *b*_0_ = 1. (Fig. 1a).

To study the evolutionary history of these *n* cells, we aim to construct a B cell lineage tree *T* that jointly describes the evolution of the DNA sequences **A** = [**a**_0_, **a**_1_, …, **a**_*n*_]^⊤^ and their isotypes **b** = [*b*_0_, *b*_1_, …, *b*_*n*_]^⊤^. As such, each node *v* of *T* will be labeled by a sequence *α*(*v*) *∈* Σ^*m*^ and isotype *β*(*v*) *∈* [*r*]. In particular, the root *v*_0_ will be labeled by *α*(*v*_0_) = **a**_0_ and *β*(*v*_0_) = *b*_0_ = 1 while the *n* leaves *L*(*T*) = {*v*_1_, …, *v*_*n*_} of *T* will be labeled by sequence *α*(*v*_*i*_) = **a**_*i*_ and isotype *β*(*b*_*i*_) = *b*_*i*_ for each *i ∈* [*n*]. A key property of isotype evolution is that it is *irreversible*. As such, the isotype *β*(*u*) of an ancestral cell *u* must be less than or equal to the isotype *β*(*v*) of its descendants *v*. More formally, we have the following definition of a B cell lineage tree.

### Definition 1.

A rooted tree *T* whose nodes are labeled by sequences *α* : *V* (*T*) → Σ^*m*^ and isotypes *β* : *V* (*T*) → [*r*] is a *B cell lineage tree* for MSA **A** = [**a**_0_, **a**_1_, …, **a**_*n*_]^⊤^ and isotypes **b** = [*b*_0_, *b*_1_, …, *b*_*n*_]^⊤^ provided (i) *T* has *n* leaves *L*(*T*) = {*v*_1_, …, *v*_*n*_} such that each leaf *v*_*i*_ *∈ L*(*T*) is labeled by sequence *α*(*v*_*i*_) = **a**_*i*_ and isotype *β*(*v*_*i*_) = *b*_*i*_, (ii) the root node *v*_0_ of *T* is labeled by sequence *α*(*v*_0_) = **a**_0_ and isotype *β*(*v*_0_) = *b*_0_, and (iii) for all nodes *u, v ∈ V* (*T*) such that *u* is ancestral to *v* it holds that *β*(*u*) ≤ *β*(*v*).

In the following, we will refer to B cell lineage trees as *lineage trees*. Lineage trees typically have shallow depth due to the limited number of mutations introduced during SHM, making parsimony a reasonable optimization criterion [12–14]. Given a lineage tree *T*, the SHM parsimony score is computed as

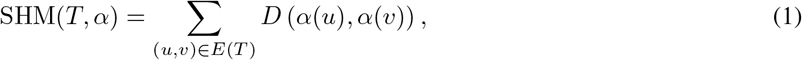

where *D* (*α*(*u*), *α*(*v*)) is the Hamming distance [22] between sequences *α*(*u*) and *α*(*v*). However, one common challenge of using parsimony to model SHM is that it often results in a large number of candidate lineage trees with equal optimal parsimony score. In addition, many inferred lineage trees contain *polytomies*, or internal nodes with out-degree greater than 2. To overcome these two challenges, we propose to infer lineage trees that optimize both sequence evolution (SHM) and isotype evolution (CSR).

Similarly to SHM, one could model the evolution of CSR using unweighted parsimony. That is, one would prefer lineage trees *T* with isotypes *β* : *V* (*T*) → [*r*] that minimize the number of isotype changes, i.e., Σ_(*u,v*)∈*E*(*T*)_ *D* (*β*(*u*), *β*(*v*)). However, there are two issues with this approach. First, it does not appropriately penalize lineage trees that violate the irreversible property of isotype evolution [13]. Second, it does not account for the fact that given an isotype starting state the probability of transitioning to each of the possible isotype states is not necessarily equal. In fact, knowing these probability distributions is useful for researchers looking to gain basic insight into the patterns and casual factors of class switch recombination [23]. Therefore, we seek to develop an appropriate evolutionary model for CSR that captures the irreversible property of class switching and models preferential isotype class transitions.

We propose a state or tree dependence model [24, 25] evolutionary model for CSR, which models the joint probability distribution of a random variable vector under Markov-like assumptions on a given tree (Appendix A). Here, the random variables of interest in this state tree model are the isotypes *β*(*v*) of each node *v* in lineage tree *T*. This model is parameterized by a probability distribution over the isotype of the root and isotype transition probabilities. As the root *v*_0_ of a lineage tree *T* is a naive B cell post V(D)J recombination, the isotype *β*(*v*_0_) is always 1 (IgM) and the probability distribution of *β*(*v*_0_) is defined as Pr(*β*(*v*_0_) = 1) = 1 and 0 otherwise. Intuitively, isotype transition probabilities captures the conditional probability of a descendant isotype given the isotype of its parent subject to irreversible isotype evolution. Next, we give a formal definition of isotype transition probabilities.

### Definition 2.

An *r* × *r* matrix **P** = [*p*_*s,t*_] is an *isotype transition probability matrix* provided for all isotypes *s, t ∈* [*r*] it holds that (i) *p*_*s,t*_ ≥ 0, (ii) *p*_*s,t*_ = 0 if *s > t*, and 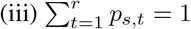 for all isotypes *s ∈ [r]*.

We define the joint likelihood CSR(*T, β*, **P**) of the observed isotypes **b** for isotype transition probabilities **P** and any lineage tree *T* whose leaves have isotypes **b** as

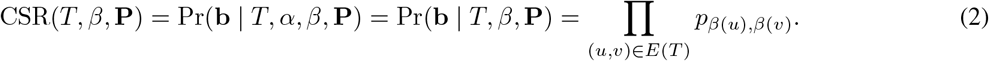

We consider the problem of simultaneously inferring a lineage tree *T* with nodes labeled by sequences *α*(*v*) and isotypes *β*(*v*) given MSA **A** and isotypes **b** that optimizes both SHM(*T, α*) and CSR(*T, β*, **P**). However, a significant barrier to solving this problem is that isotype transition probabilities **P** are unknown and need to be inferred. While there have been experimental studies that estimate these quantities under specific biological conditions [23], there currently exists no computational methods to infer these probabilities directly from a sequencing experiment. Here, we will leverage that typical single-cell sequencing experiments yield data from a set of diverse B cell lineages, where each lineage is commonly referred to as a *clonotype*. A common pre-processing step is to first cluster the *n* sequenced cells into *k* clonotypes, where each clonotype *j* contains *n*_*j*_ cells (Fig. 1c). We follow convention and group B cells based on shared V(D)J alleles in heavy and light chains into a clonotype. Moreover, we reason that under many experimental conditions, the transition probabilities between isotypes will be similar to those within clonotypes. Thus, inferring these isotype transition probabilities for *k* clonotypes will yield higher accuracy than inferring isotype transition probabilities for a single lineage in isolation. This leads to the following problem statement.

### Problem 1

(B CELL Lineage Forest Inference (BLFI)). Given MSAs **A**_1_, …, **A**_*k*_ and isotypes **b**_1_, …, **b**_*k*_ for *k* clonotypes, find isotype transition probabilities **P**^*^ for *r* isotypes and lineage trees 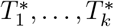 for (**A**_1_, **b**_1_), …, (**A**_*k*_, **b**_*k*_) whose nodes are labeled by sequences 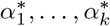 and isotypes 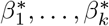, respectively, such that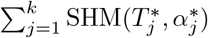 is minimum and then 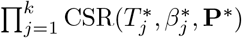 is maximum.

In other words, we prioritize the SHM objective and then among all solutions with minimum SHM score we prefer those that additionally maximize the CSR objective. It is easy to see that the BLFI problem is NP-hard via a simple reduction from the Large Parsimony problem — see Appendix B.1 for more details.

## 3 Methods

We introduce Tree Inference of B cell Clonal Lineages, or TRIBAL, a method to solve the BLFI problem. This method is based on two key ideas that allow us to effectively solve the BLFI problem.

First, the lexicographical ordering of our two objectives — optimizing for SHM followed by CSR — enables one to use the following two-stage approach (Fig. 1c). In the first stage, we use existing maximum parsimony methods to generate a set 𝒯 of input trees — also called a *maximum parsimony forest* — for each clonotype such that each tree *T ∈* 𝒯 minimizes the objective SHM(*T, α*). To do so, we provide these methods only the sequence information **A** to enumerate a solution space 𝒯 of trees whose nodes are labeled by sequences *α*_1_, …, *α*_| *𝒯* |_. In the second stage, we incorporate isotype information **b** to further operate on the set 𝒯 and additionally optimize CSR(*T, β*, **P**) in such a manner that maintains optimality of the SHM objective. We note that a lexicographically optimal lineage tree *T*^*^ does not necessarily need to be an element of 𝒯, but instead it suffices that the evolutionary relationships in tree *T*^*^ are a refinement of the evolutionary relationships described by some tree *T* among the set 𝒯 of input trees. More specifically, a *refinement T* ′ of tree *T* is obtained by a series of expand operations such that an expand operation of node *v* consists of splitting node *v* into *v* and *v*′, joining them with an edge (*v, v*′) and then redistributing the children of *v* to be descendants of either *v* or *v*′. We have the following key proposition and corollary.

### Proposition 1.

For any tree *T* labeled by sequences *α* and refinement *T* ′ of *T*, there exists a sequence labeling *α*′ for *T* ′ such that SHM(*T, α*) = SHM(*T*′, *α*′).

*Proof*. The sequencing labeling *α*′ is found by setting *α*′(*v*) = *α*(*v*) and *α*′(*v*′) = *α*(*v*) during each EXPAND operation. By construction, the new edge (*v, v*′) has *D*(*α*′(*v*), *α*′(*v*′)) = 0 and every original edge maintains its original Hamming distance in *T* ′. Therefore, SHM(*T, α*) = SHM(*T*′, *α*′).

### Corollary 1.

Any lineage tree *T* ′ that lexicographically optimizes SHM(*T*′, *α*′) and then CSR(*T*′, *β*′, **P**) must be a refinement of some tree *T* optimizing only SHM(*T, α*).

Therefore, our sought lineage tree *T*^*^ that lexicographically optimizes both objectives must be a refinement of some tree *T* in the set 𝒯.

The second key idea is that the inference of optimal lineage trees 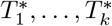 is conditionally independent when given isotype transition probabilities **P**. This motivates the use of a coordinate ascent algorithm where we randomly initialize isotype transition probabilies **P**^(1)^ (Sec. 3.1 and Fig. 2a). Then, at each iteration ℓ, we use isotype transition probabilities **P**^(ℓ)^ and the input set 𝒯_*j*_ of trees to independently infer an optimal lineage tree 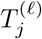 for each clonotype *j* (Sec. 3.2 and Fig. 2b). This is then followed by estimating updated isotype transition probabilities **P**^(ℓ+1)^ given trees 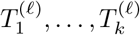 (Sec. 3.3 and Fig. 2c). We terminate upon convergence of our CSR objective or when exceeding a specified number of maximum iterations.

**Figure 2.**
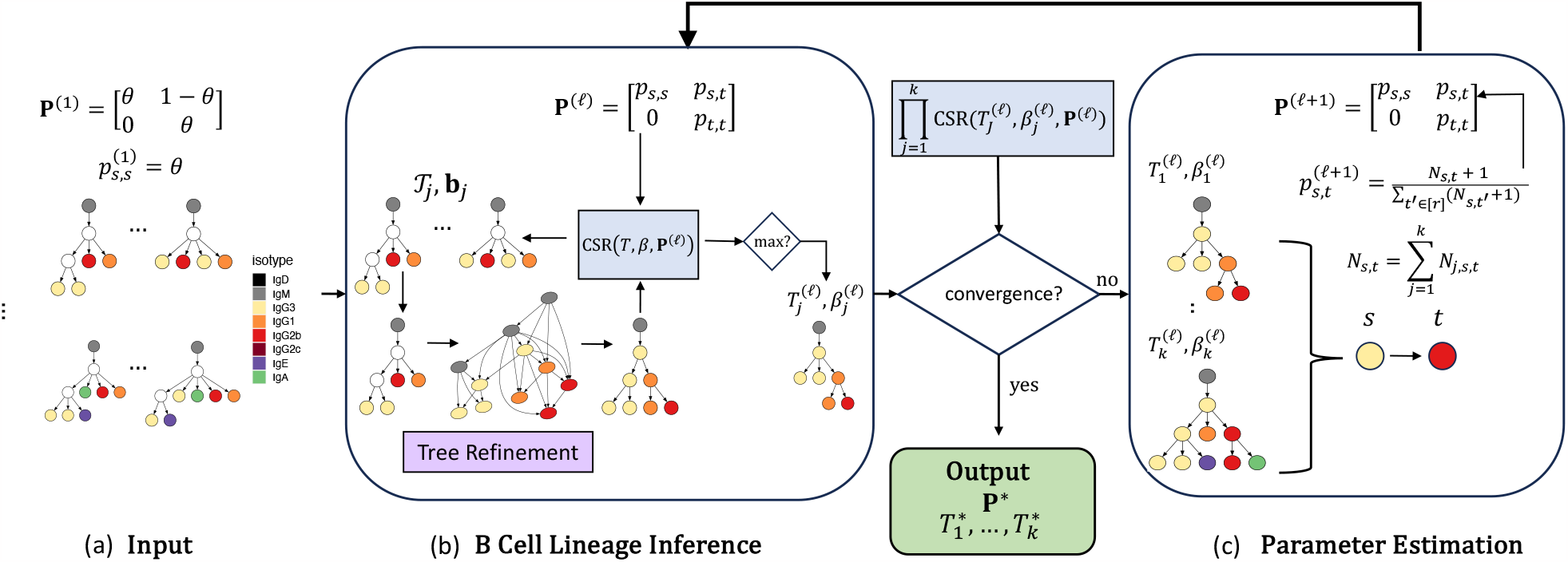
TRIBAL infers B cell lineage forest 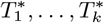 and isotype transitions P*\** for *k* clonotypes utilizing coordinate ascent. (a) The inputs to TRIBAL are isotype transition probabilities **P**^(1)^, which are initialized given a parameter *θ* ∈ [0.5, 1], and a tuples (𝒯_1_, **b**_1_) …, (𝒯_*k*_, **b**_*k*_), where set 𝒯_*j*_ are maximum parsimony trees for MSA **A**_*j*_ and **b**_*j*_ are the observed isotypes of the *nj* cells of clonotype *j*. (b) Conditioning on isotype transition probabilities **P**^(*ℓ*)^, a B cell lineage tree 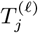 with nodes labeled by isotypes 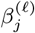 is inferred for each clonotype *j* by solving the MPTR problem for each tree in the input set 𝒯 *j*. (c) Convergence between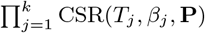 for iterations *ℓ* and *ℓ −* 1 is checked. If the difference has not converged, isotype transition probabilities **P**^(*ℓ*+1)^ are updated using maximum likelihood estimation. If the difference has converged, the current inferred B cell lineage forest and isotype transition probabilities **P** are output. Multiple restarts may be performed for different values of *θ*.

TRIBAL is implemented in Python 3, is open source (BSD-3-Clause license), and is available at https://github.com/elkebir-group/tribal.

### 3.1 Input

The input to TRIBAL is a set of *k* clonotypes with corresponding maximum parsimony forest 𝒯_*j*_ for each clonotype 𝒯 *j*. In addition, we are given isotypes **b**_*j*_ labeling the leaves of trees 𝒯_*j*_ for each clonotype *j* (Fig.1c, Fig. 2a). Obtaining this input requires a number of preprocessing steps of a scRNA-seq dataset (Fig. 1), including (i) BCR assembly and isotype calling of each sequenced cell, (ii) clonotyping or clustering the cells based on a shared germline alleles for both the heavy and light chains, (iii) obtaining an MSA for sequences within a clonotype and (iv) finding a parsimony forest for each MSA of a clonotype. There preprocessing steps are not part of TRIBAL.

For our coordinate ascent approach, we also require an initialization for the isotype transition probabilities (Fig. 2a). We set the initial transition probabilities to reflect the observation that under baseline conditions, the probability of a B cell undergoing class switching is lower than the probability of it maintaining its original antibody class. [4, 23]. Thus, we initialize **P**^(1)^ such that *p*_*s,s*_ *> p*_*s,t*_ for all isotypes *s* and *t*. Let *θ* ∈ [0.5, 1] be the probability that a B cell does not class switch, i.e., *p*_*s,s*_ = *θ* for each isotype *s* < *r* and *p*_*s,s*_ = 1 if *s* = *r*. We enforce irreversibility such that *p*_*s,t*_ = 0, if *s > t*. We then initialize the remaining parameters uniformly, i.e., *p*_*s,t*_ = (1 − *p*_*s,s*_)*/*(*r* − *s*) where *r* is the total of number isotypes. We conduct multiple restarts, varying *θ* ∈ [0.5, 1] in each restart.

### 3.2 B cell lineage tree inference via refinement

As described above, the inference of optimal lineage trees 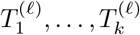 is conditionally independent given isotype transition probabilities **P**^(ℓ)^. We therefore focus our discussion on how TRIBAL infers a B cell lineage tree 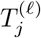 for a single clonotype *j* during iteration ℓ given isotype transition probabilities **P**^(ℓ)^. By Corollary 1, we solve this problem by finding an optimal refinement *T* ′ and corresponding isotype labeling *β*′ for each tree *T* in the input set 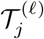 and select the one that maximizes our CSR objective (Fig. 2b). Maximizing the log-likelihood of CSR(*T, β*, **P**) is equivalent to maximizing a weighted parsimony criterion. This leads to the following problem statement.

#### Problem 2

(Most Parsimonious Tree Refinement (MPTR)). Given a tree *T* on *n* leaves, isotypes **b** = [*b*_0_, …, *b*_*n*_] and isotype transition probabilities **P**, find a tree *T* ′ with root 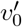 and isotype labels *β*′ : *V* (*T* ′) → [*r*] such that (i) *T* ′ is a refinement of *T*, 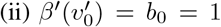, 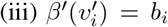 for each leaf 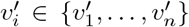 and (iv) log CSR(*T*′, *β*′, **P**) is maximum.

We prove in Appendix B.1 that the MPTR problem is NP-hard. Therefore, we convert this problem to a graph problem (Fig. S2). For each node *u* in tree *T*, multiple copies (*u, s*) are added to the expansion graph *G*_*T*,**b**_, one for each possible isotype *s* ∈ [*r*] subject to additional constraints depending on the observed isotypes of the leaves descendant from *u*. The edges ((*u, s*), (*v, t*)) in this graph have weight log *p*_*s,t*_. We then solve the MPTR problem by finding a valid, maximum weight subtree (*T* ′, *β*′) in the expansion graph *G*_*T*,**b**_ such that *T* ′ with isotype labels *β*′ is a most parsimonious refinement of *T*. This is achieved via a mixed-integer linear programming (MILP) formulation, similar to that used to solve the Steiner minimal tree problem [26]. See Appendix C.1 for more details on the construction of the expansion graph from tree *T* and leaf isotypes **b**, proof of correctness for this algorithm and the MILP formulation.

To obtain outputs 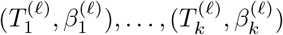, we set 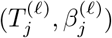 to the tree *T*′ and corresponding isotype labeling *β*′ that maximize CSR(*T*′, *β*′, **P**^(ℓ)^) among all input trees 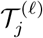 for each clonotype *j* ∈ [*k*].

### 3.3 Parameter estimation

In the next step, given tuples 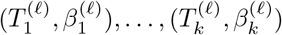 of lineage trees and isotype labels for iteration ℓ, we seek updated isotype transition probabilities **P**^(ℓ+1)^ for iteration ℓ + 1 such that 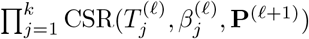 is maximum (Fig. 2c). This is achieved via maximum likelihood estimation. For fixed lineage trees *T*_1_, … *T*_*k*_ and isotypes *β*_1_, …, *β*_*k*_, our CSR objective can be rewritten as

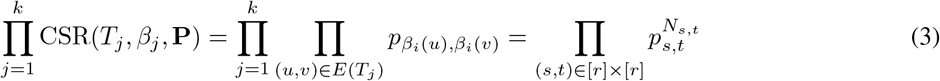

where 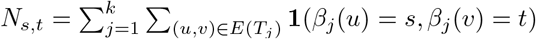 is the number of transition from *s* to *t* across all selected trees and clonotypes. Thus, we seek isotype transition probabilities **P** that maximize (3) subject to the constraints that 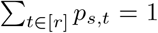 for every isotype *s*. This constrained optimization problem is easily solved with the aid of Lagrange multipliers (see Appendix C.2 for details). Additionally, we add pseudocounts of 1 to avoid overfitting the data in the event of unobserved transitions, yielding the following updated probabilities 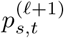 for iteration ℓ + 1.

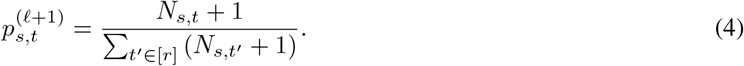

After estimating **P**^(ℓ+1)^, we then infer 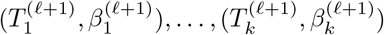, as discussed in Section 3.2.

## 4 Results

We evaluated TRIBAL on both *in silico* experiments (Sec. 4.1) and two sets of experimental scRNA-seq data. The first assessed performance on a model immunological system for affinity maturation (Sec. 4.2) and the second on a study of the relationship between age-associated B cells and autoimmune disorders (Sec. 4.3).

### 4.1 In silico experiments

We designed *in silico* experiments to evaluate TRIBAL with known ground-truth isotype transition probabilities **P** and lineage trees *T* labeled by sequences *α* and isotypes *β*. Specifically, we used an existing BCR phylogenetic simulator [13] that models SHM but not CSR. We generated isotype transition probabilities **P** with *r* = 7 isotypes (as in mice) under two different models of CSR. Briefly, both CSR models assume the probability of not transitioning is higher than the probability of transitioning, but in the *sequential model* there is clear preference for transitions to the next contiguous isotype, while in the *direct model* the probabilities of contiguous and non-contiguous class are similar. Given **P**, we evolved isotype characters down each ground truth lineage tree *T*. We generated 5 replications of each CSR model for *k* = 75 clonotypes and *n* ∈ { 35, 65 } cells per clonotype, resulting in 20 *in silico* experiments, yielding a total of 1500 ground truth lineage trees. In addition to comparing TRIBAL to existing methods including dnapars [7], dnaml [7] and IgPhyML [10], we also compared to a version of TRIBAL without tree refinement, denoted as TRIBAL-NO REFINEMENT (TRIBAL-NR). To obtain the input set 𝒯 _*j*_ of trees with maximum parsimony for each clonotype *j*, we utilized dnapars [7]. We refer to Appendix D for additional details on the simulations. In the following we focus our discussion on *in silico* experiments with *n* = 35 cells per clonotype (see Fig. S5 for *n* = 65).

To evaluate accuracy of isotype transition probability inference, we used *Kullback–Leibler (KL) divergence* [27] to compare the inferred transition probability distribution 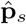 of each isotype *s* to the simulated ground truth distribution **p**_*s*_. KL divergence is defined as 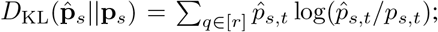 the lower the KL divergence, the more similar the two distributions. Since no existing methods infer isotype transition probabilities, we restricted this analysis to TRIBAL and TRIBAL-NR. Overall, we observed good concordance between simulated and TRIBAL inferred isotype transition probabilities (Fig. 3a). Specifically, TRIBAL had lower median KL divergence than TRIBAL-NR for all isotype starting states, except IgA, which is trivially 0, under both direct and sequential CSR models (direct: median of 0.15 vs. 0.73; sequential: median of 0.099 vs. 0.55). We observed improved performance of TRIBAL (but not for TRIBAL-NR) for *n* = 65 cells per clonotype (Fig. S5a and Fig. S6). To assess the sensitivity of TRIBAL to infer isotype transition probabilities with fewer than *k* = 75 clonotypes, we downsampled the 75 clonotypes to 25 and 50 clonotypes per experiment. We observed similar trends for experiments with *k∈* {25, 50} clonotypes, with TRIBAL continuing to outperform TRIBAL-NR while still achieving small KL divergences even as *k* decreases (Fig. S7). These findings demonstrate that tree refinement is key to accurately estimate isotype transition probabilities.

**Figure 3.**
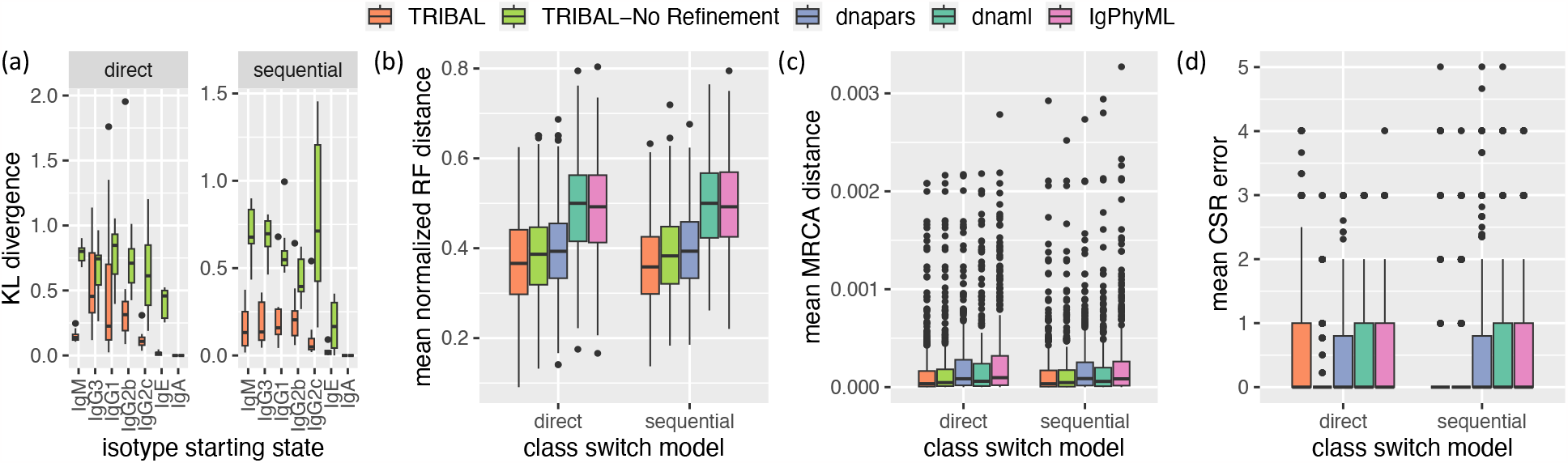
TRIBAL accurately infers isotype transition probabilities on simulated data while outperforming existing methods on lineage tree inference. Simulation results shown are for 5 replications with *k* = 75 clonotypes per replication and *n* = 35 cells per clonotype. (a) KL divergence between inferred isotype transition probabilities and the reference ground truth distribution. (b) mean Robinson-Foulds distance between ground truth and inferred lineages tree per clonotype. (c) mean MRCA distance (26) between ground truth and inferred lineage trees per clonotype. (d) mean CSR error between ground truth and inferred B cells.

Next, we assessed the accuracy of lineage tree inference using the *Robinson-Foulds (RF) distance* [28] normalized by the total number of bipartitions in the ground truth *T* and inferred lineage tree 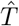 (25). Since TRIBAL, TRIBAL-NR and dnapars return multiple optimal solutions, we report the mean of the lineage tree inference metrics over all optimal solutions. We found TRIBAL had the lowest mean normalized RF distance for both direct and sequential CSR models (Fig. 3b). Overall, dnaml (median: 0.5) and IgPhyML (median: 0.49) had the worst performance on normalized RF. Interestingly, even though the starting trees of dnapars are used by TRIBAL, both TRIBAL (median: 0.36) and TRIBAL-NR (median: 0.38) outperformed dnapars (median: 0.39), showing the importance of using isotype information to resolve phylogenetic uncertainty.

While normalized RF distance only assesses the accuracy of the tree topology, it is important to also assess the accuracy of the ancestral sequence reconstruction. To that end, we used a metric called *Most Recent Common Ancestor (MRCA) distance* (26) introduced by Davidsen and Matsen [13]. For any two simulated B cells (leaves), the MRCA distance is the Hamming distance between the MRCA sequences of these two B cells in both the ground truth and inferred lineage trees. This distance is then averaged over all pairs of simulated B cells — see Appendix D.4 and Fig. S4a for additional details. Again, we report the mean of over all optimal solutions for TRIBAL, TRIBAL-NR and dnapars. We found TRIBAL outperformed all other methods (Fig. 3c), achieving the lowest overall median MRCA distance (3.46 *×* 10^−5^), followed by TRIBAL-NR (4.63 *×* 10^−5^). IgPhyML had the worst performance with a median of 8.78 × 10^−5^. Performance trends were consistent between methods across both CSR models.

Lastly, we assessed the accuracy of isotype inference by a new metric called *CSR error*, which is computed for each B cell *i* and clonotype *j* and is the absolute difference between the number of ground-truth class switches and inferred number of class switches that occurred along its evolutionary path from the root — see Appendix D and Fig S4b for additional details. Since dnaml, dnapars and IgPhyML do not infer isotypes for internal nodes, we pair these methods with the Sankoff algorithm [29] using *w*_*s,t*_ equals 1 if *s* = *t*, 0 if *s* < *t* and ∞ otherwise. We account for the presence of multiple solutions by taking the mean across solutions. All methods had a median CSR error of 0 for both the direct and sequential models (Fig. 3d). Therefore, we utilized the third quartile for a more robust comparison. We found that under the direct model TRIBAL-NR (third quartile: 0) was the best performing method, while dnapars was second best (third quartile 0.8) and all other methods, including TRIBAL had a third quartile of 1. We observed a slight tendency of TRIBAL to overestimate the number of transitions due to the tree refinement step, while other methods tended to underestimate the number of transitions. However, under a sequential model, where refinement is helpful in accurately capturing sequential state transitions, we found that TRIBAL was tied with TRIBAL-NR for the best performance (third quartile: 0). All other methods had similar performance between both CSR models for this metric.

We observed similar trends on these metrics for *in silico* experiments containing *n* = 65 cells per clonotype for *k* = 75 clonotypes (Fig. S5). In summary, these results suggests that the incorporation of isotype data and the use of tree refinement are beneficial for both lineage tree inference and ancestral sequence and class switch reconstruction.

### 4.2 Keyhole limpet haemocyanin (NP-KLH) antigen immune response studies

We applied TRIBAL as well as IgPhyML to 10 × 5′ scRNA-seq data of B cells extracted from mice immunized with nucleoprotein keyhole limpet haemocyanin (NP-KLH), a commonly used antigen in the study of antibody affinity maturation [30]. Our goal was to determine whether these methods recapitulate known patterns of B cell lineage evolution for this well-studied antigen using data from two studies and to compare the lineage trees inferred by each method. The first dataset (NP-KLH-1) was generated from C57BL/6 mice that were immunized with NP-KLH and total germinal center B cells were extracted 14 days after immunization [20]. The other two datasets came from a single study in which C57BL/6 mice were immunized with NP-KLH (NP-KLH-2a and NP-KLH-2b) and NP-specific germinal center B cells were extracted 13 days after immunization [31]. We utilized the standard 10 × Cell Ranger [21] single-cell bioinformatics pipeline to generate sequence **a**_*i*_ and isotype *b*_*i*_ for each cell *i*. We used Dandelion [32] to remove doublets, reassign alleles, and cluster the cells into clonotypes. We identified clonotype MSAs **A**_1_, …, **A**_*k*_ based on shared V(D)J alleles for the heavy chain using the dowser package [14]. Finally, we excluded clonotypes with fewer than 5 cells. This yielded a total of *n* = 2670 sequenced B cells clustered into *k* = 295 clonotypes. We exclude methods that rely on sequence abundance as a key signal, such as GCTree [12] and ClonalTree [17] as we observed very few duplicated sequences within each clonotype. Fig. 4a shows the distribution of isotypes by dataset and Table S1 includes a more detailed summary of each dataset.

**Figure 4.**
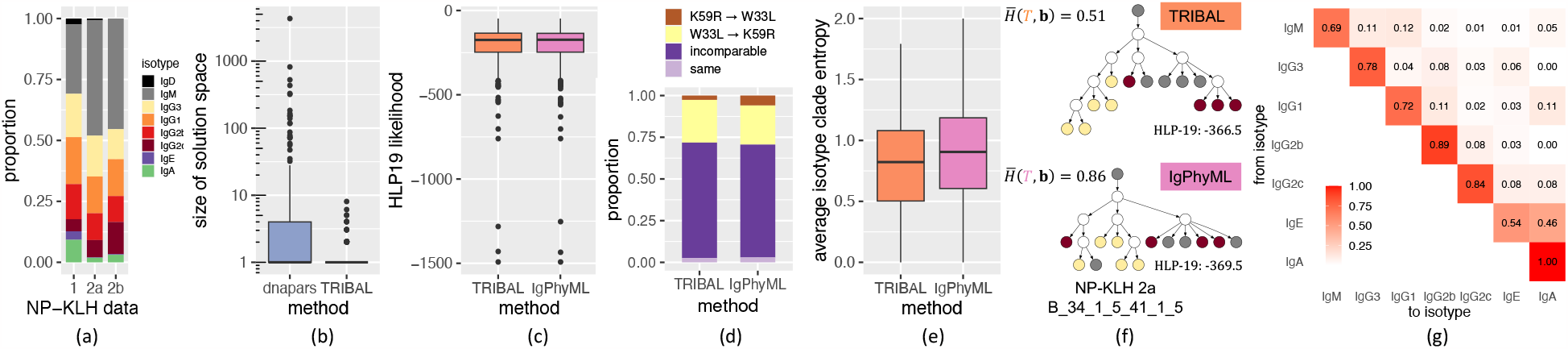
Comparison between TRIBAL and IgPhyML on the NP-KLH data. (a) The distribution of isotypes **b** in each dataset. (b) A comparison of the solution space of dnapars versus TRIBAL. (c) The distribution of the HLP19 codon-substitution likelihood [11] for lineage trees inferred by TRIBAL and IgPhyML. (d) Observed distribution of evolutionary relationships between the W33L and K59 in clonotypes where both mutations are present. (e) The distribution of the average clade entropy with respect to an isotype labeling **b** of the leafset. (f) A comparison of lineage trees inferred for clonotype NP-KLH-2a B 34 1 5 41 1 5 with the average isotype clade entropy 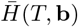 reported for each inferred tree. (g) TRIBAL inferred isotype transition probabilities **P** for NP-KLH-1.

We used dnapars [7] to infer TRIBAL’s input set 𝒯_*j*_ for each clonotype *j*. We found that TRIBAL’s use of isotype information significantly reduced the number of optimal solutions identified by dnapars (mean: 31.5 vs. 1.3, max: 4310 vs. 8) — see Fig 4b. While IgPhyML, a maximum likelihood method using the HLP19 codon-substitution model [10, 11], infers only a single tree per clonotype, it is important to note that there might be multiple trees with maximum likelihood in the solution space. Indeed, we found high concordance of HLP19 likelihoods between the TRIBAL and IgPhyML inferred lineage trees, with a small overall mean absolute deviation of 0.97 (Fig. 4c, Fig. S8). We even observed that TRIBAL had a greater likelihood than IgPhyML in 59.3% of the clonotypes. Thus, TRIBAL resulted in a significant reduction in the size of the solution space compared to the maximum parsimony method dnapars with similar (and sometimes better) HLP19 likelikelihood as IgPhyML, illustrating how isotype information can be used to effectively reduce phylogenetic uncertainty.

Next, we assessed whether the lineage trees inferred by TRIBAL recapitulated expected biological trends for the NP-KLH model system. Previous research has indicated that the IGHV1-72*01 (VH186.2) variable gene in the heavy chain is preferentially used in the anti-NP response in C57BL/6 mice [33] via two distinct mutation paths: one in which a W33L mutation greatly increases affinity to NP, and another in which high affinity is achieved through K59R and the accumulation of several other mutations, such as S66N [34, 35]. Consequently, we expect that for clonotypes containing both W33L and K59R, these mutations would tend towards occurring in distinct lineages of the tree. We analyzed the inferred pairwise relationships between W33L & K59R in the 26 lineage trees that contained both mutations. Specifically, we categorized the relationship as K59R→ W33L if K59R was ancestral to W33L, W33L→ K59R if W33L was ancestral to K59R, *incomparable* if W33L and K59R occurred on distinct lineages of the tree and *same* if they were introduced on the same edge of the lineage tree. Indeed, we confirmed the tendency for mutual exclusivity of W33L and K59R by finding that the proportion of pairwise introductions categorized as *incomparable* was 0.69 and 0.67 for TRIBAL and IgPhyML, respectively (Fig. 4d). Additionally, it has been suggested that W33L mutations appear relatively early during the anti-NP response, whereas the K59R and S66N mutations typically appear later in the evolutionary history [36]. Defining level as the length of the shortest path from the MRCA of all B cells, we observed that W33L occurred at a median level of 1 for both TRIBAL and IgPhyML while both the K59R and S66N mutation occurred at a median level 2 for both methods (Fig. S9). This indicated that W33L was typically introduced earlier in the evolutionary history of a clonotype than K59R and S66N. Thus, both TRIBAL and IgPhyML trees recapitulate expected mutation patterns for this model system.

We next assessed the extent of agreement with isotype information. While TRIBAL infers isotype labels of ancestral nodes, IgPhyML does not have this capability. Therefore, we developed a new metric called *average isotype clade entropy*, which is computed with respect to the isotype labeling of the leaf set. For this metric, we compute the entropy of clade *u* in tree *T* with respect to all isotype leaf labels that are descendants of node *u*, taking the average entropy over all non-trivial clades, which excludes the root and the leaves (Appendix E.1). As IgPhyML returns bifurcating trees, we collapse edges with zero branch length for a more fair comparison of this metric. We observed lower average isotype clade entropy for TRIBAL (median: 0.82) versus IgPhyML (median 0.91) 4f. Fig. 4e depicts the lineage tree inferred by TRIBAL and IgPhyML for the NP-KLH-2a dataset (clonotype B 34 1 5 41 1 5). The TRIBAL inferred tree for this clonotype had lower isotype clade entropy than IgPhyML (TRIBAL: 0.51 vs IgPhyML: 0.86) while also resulting in a greater HLP19 likelihood (TRIBAL: −366.5 vs IgPhyML: −369.5). Thus, we find that the trees identified by TRIBAL are in better agreement with the leaf isotypes than IgPhyML.

In addition to the inferred B cell lineage trees, TRIBAL also inferred isotype transition probabilities **P** for each dataset (Fig. 4g, Fig S10). All three inferred isotype transition probability matrices more closely matched a CSR model of direct switching as opposed to a strictly sequential model. To compare the consistency of these estimates across datasets, we computed the Jensen-Shannon divergence (JSD) between the distribution of isotype transition probabilities for each isotype starting state IgM through IgG2c for each dataset pair. We observed low JSD (median: 0.029) across a total of 15 pairwise comparisons, suggesting consistent estimates between isotype transition probabilities.

In summary, these analyses show that the inclusion of isotype information and tree refinement has the potential to yield high quality lineage tree inference, even under a simpler model of SHM, i.e., parsimony. Moreover, the TRIBAL inferred lineage trees additionally optimize an evolutionary model of CSR, yielding lower isotype entropy partitions of the leaf set than IgPhyML. Finally, the additional inference of isotype transitions probabilities **P** has the potential to distinguish between direct versus sequential switching events.

### 4.3 Age-associated B cell (ABC) datasets

We evaluated TRIBAL on three scRNA-seq datasets with V region sequencing that investigated the relationship between age-associated B cells (ABCs) and autoimmune disorders [37]. For each dataset, B cells were extracted from the spleen of a MRL/lpr female mouse and sequenced using 10 *×*5′ scRNA-seq. The data was processed by the 10*×* Cell Ranger [21] single-cell bioinformatics pipeline to generate sequence **a**_*i*_ and isotype *b*_*i*_ for each cell *i*. Nickerson et al. [37] identified clonotype MSAs **A**_1_, …, **A**_*k*_ based on shared V(D)J alleles for the heavy chain using the dowser package [14] and inferred B cell lineage trees using IgPhyML for each clonotype. After filtering out clonotypes with fewer than 5 sequences, we retained 599 B cells and 54 clonotypes across the three datasets (Table S2). Fig. 5a shows the proportion of isotypes and annotations by mouse for the retained B cells. Of these 54 clonotypes, 35 had more than one distinct isotype across the sequenced B cells, with a median of 3 distinct isotypes per clonotype.

**Figure 5.**
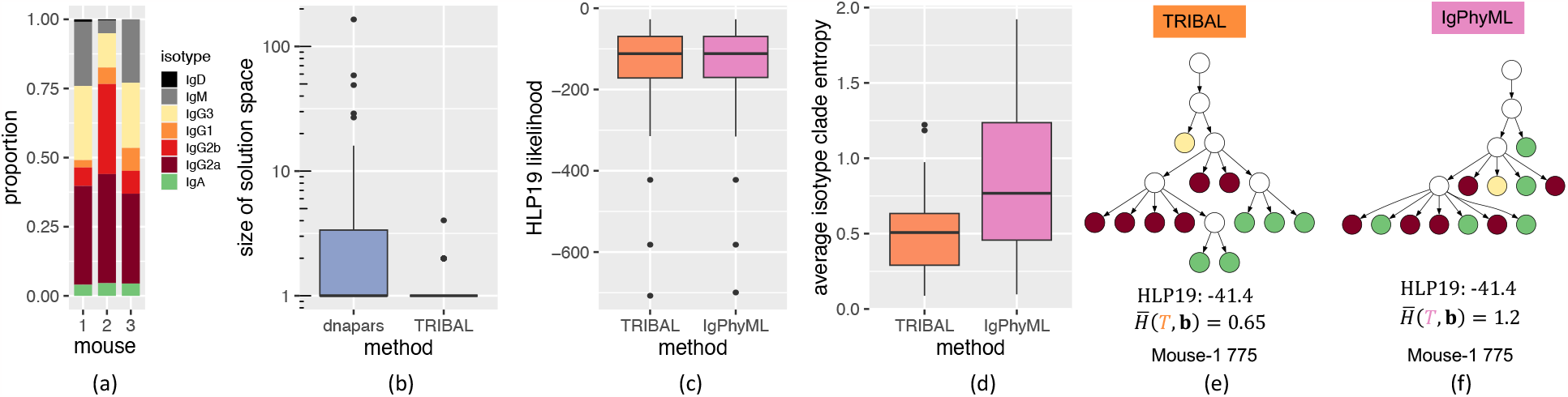
Comparison between TRIBAL and IgPhyML on ABC data. (a) Distribution of B cell isotypes. (b) A comparison of the solution space of dnapars versus TRIBAL. (c) The distribution of the HLP19 codon-substitution likelihood [11] for lineage trees inferred by TRIBAL and IgPhyML. (d) Comparison of average isotype clade entropy for TRIBAL versus IgPhyML. (e,f) Comparison of inferred B cell lineage trees by TRIBAL (e) and IgPhyML (f) for clonotype Mouse-1 775 — see (a) for color legend.

We ran TRIBAL separately on each of the three mouse datasets, obtaining a maximum parsimony forest 𝒯_*j*_ for each clonotype *j* via dnapars. Similar to our NP-KLH analysis, we found that TRIBAL effectively utilized the additional isotype data to reduce the number of optimal solutions identified by dnapars (mean: 8.1 vs. 1.3, max: 165 vs. 4) — see Fig. 5b. The HLP19 likelihood of the TRIBAL inferred lineage trees had high concordance with the IgPhyML inferred trees (mean absolute deviation: 0.97), with TRIBAL yielding a higher likelihood for 53% of the clonotypes (Fig. 5c, Fig. S11). The average isotype clade entropy for the 35 clonotypes with more than one distinct isotype was significantly lower for TRIBAL than for IgPhyML (median: 0.49 vs. 0.77) — see Fig. 5d. An example comparison is shown in Fig. 5e,f for clonotype Mouse-1 775. The tree refinement step of TRIBAL yielded a tree with a significantly lower average isotype clade entropy when compared to IgPhyML (0.65 vs. 1.2) while both trees had identical HLP19 likelihoods (− 41.4). Finally, we observed that the isotype transition probabilities reveal evidence for both direct and sequential switching of isotypes (Fig. S12).

In summary, both TRIBAL and IgPhyML yield lineage trees with very similar HLP19 likelihoods, giving support for the validity of the TRIBAL inferred lineage trees in terms of sequence evolution. However, TRIBAL jointly optimizes evolutionary models for both SHM and CSR, yielding trees with lower average isotype clade entropy.

## 5 Conclusion

The development and application of methods for inferring B cell lineage trees and isotype transition probabilities from scRNA-seq data is crucial for advancing our understanding of the immune system. In this work, we introduced TRIBAL, a method to infer B cell lineage trees and isotype transition probabilities from scRNA-seq data. TRIBAL makes use of existing maximum parsimony methods to optimize an evolutionary model for SHM, then incorporates isotype data to find the most parsimonious refinement, i.e., maximizing the CSR likelihood, among the input set of trees. We proved that the subproblem of finding a refinement maximizing the CSR likelihood is NP-hard. We demonstrated the effectiveness of TRIBAL via *in silico* experiments and on experimental data. On *in silico* experiments, we showed the importance of tree refinement for both accurately estimating isotype transition probabilities and lineage tree inference. Furthermore, we demonstrated on experimental data that TRIBAL returns lineage trees that have similar HLP19 likelihoods, despite utilizing a less complex model for sequence evolution but yield a reduction in the entropy of the isotype leaf labelings.

There are several directions for future research. First, integration of germline “sterile” transcripts may offer a way to initialize the TRIBAL inferred isotype transition probabilities [38]. Second, many existing B cell lineage inference methods, such as IgPhyML, yield multifurcating trees when zero length branches are collapsed. There exists an opportunity to combine likelihood or distance based inference methods with the tree refinement step of TRIBAL. Third, the MPTR problem has a more general formulation with the potential for wider applications beyond the problem of B cell lineage inference. For example, sample location is useful in refining tumor phylogeny with polytomies [15]. On a related note, we hypothesize that there are special cases of the MPTR problem and its more general formulation that are in P. Such special cases may include a weight matrix with unit costs and an upper triangular weight matrix that adheres to the triangle inequality. Fourth, the assumption that a single isotype transition probability matrix is shared by all clonotypes could be relaxed to allow the inference of multiple matrices per experiment and an assignment of clonotypes to an inferred matrix. Fifth, TRIBAL could be extended to allow for the correction of inaccurately clonotyped B cells. Finally, more robust evolutionary models for SHM are needed to capture the presence of complex mutations, such as indels, introduced during affinity maturation [39, 40].

## Acknowledgments

Authors thank Harinder Singh, Ken Hoehn, Mark Shlomchik and Margie Ackerman for insightful discussions. This work was partially supported by NIH grant DP2AI177884 (A.A.K.) and by the National Science Foundation grant CCF-2046488 (M.E-K.). This work used resources, services, and support provided via the Greg Gulick Honorary Research Award Opportunity supported by a gift from Amazon Web Services.

## A Evolutionary model for class switch recombination

A dependence tree 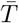 with *n* nodes, sometimes referred to as a state tree, is a tree that defines the conditional independence structure of the random variables associated with the nodes of the tree [24, 25]. Simply put, it is a type of Bayesian network, where the underlying directed acyclic graph is a tree. For each node *i* in dependence tree 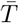, we associate a random variable *Y*_*i*_*∈ S*, where *S* is a discrete state space. Additionally, *Y*_1_ is the random variable associated with the root node. The joint probability **Y** = (*Y*_1_, …, *Y*_*n*_) of these random variables given the underlying structure of dependence tree 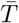 is defined as follows

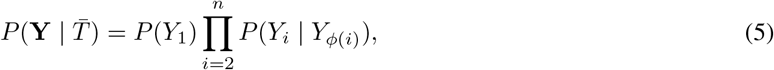

where [*n*] = {1, …, *n*} and *ϕ*(*i*) is a function that returns the parent of node *i* specified by dependence tree 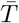. Like Markov chains, this model is parameterized by a distribution over the starting state, *π*_*s*_ = *P* (*Y*_1_ = *s*), where ∑_*i*∈[*r*]_ *π*_*s*_ = 1, and transition probabilities *p*_*s,t*_ = *P* (*Y*_*i*_ = *t* | *Y*_*ϕ*_ (*i*) = *s*) from state *s* to *t*. Transition probabilities **P** = [*p*_*s,t*_] *∈* [0, 1]^| *𝒮*|*×*| *𝒮*|^ have the property that ∑_*t∈S*_ *p*_*s,t*_ for every state *s*.

We model class switch recombination with a dependence tree 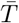 for each lineage tree *T* with nodes 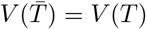 and edges 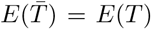 and isotype labels *β*(*v*) = *Y*_*v*_ for every node *v*. In words, the corresponding dependence tree 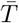 for B cell lineage tree *T* has the same topology but the dependence tree 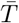 has a random variable for isotype associated with each node.

For the leaf nodes *v*_*i*_ *∈ L*(*T*), i.e., the sequenced B cells, we observe isotype *b*_*i*_ directly from scRNA-seq data, i.e., 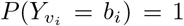. Additionally, root node *v*_0_ in *T*, represents the naive B cell post V(D)J recombination and therefore 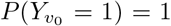, meaning that root of the dependence tree has isotype state IgM and *π*_1_ = 1. This means that any dependence tree without 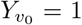 has zero probability and we omit the initial state probability term. For any other node *u* in *T*, we have *β*(*u*) = *Y*_*u*_. Thus, to compute (5) for isotype labels *β*(*v*), it suffices to know the isotype transition probabilities **P**, which we formally define below.

**(Main Text) Definition 2**. An *r* × *r* matrix **P** = [*p*_*s,t*_] is an *isotype transition probability matrix* provided for all isotypes *s, t* ∈ [*r*] it holds that (i) *p*_*s,t*_ = 0 if *t > s*, (ii) *p*_*s,t*_ ≥ 0, and (iii)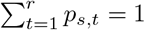 for all isotypes *s* ∈ [r].

The additional conditions on the transition probabilities beyond row stochasticity on the transition probabilities are to properly model the irreversibility of class switch recombination. Using the above model for class switch recombination, we compute the likelihood CSR(*T, β*, **P**) for observed isotypes **b** given a lineage tree with isotypes *β* and isotype transition probabilities as follows.

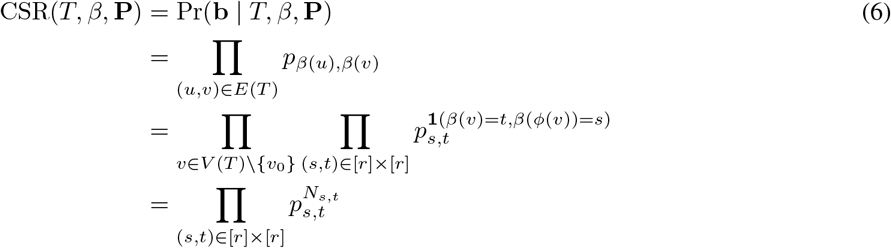

where *N*_*s,t*_ is the count of occurrences in lineage tree *T* such that *β*(*v*) = *t* and *β*(*ϕ*(*v*)) = *s*. This is easily extended for a forest of *k* lineage trees *T*_1_, …, *T*_*k*_ with corresponding isotypes *β*_1_, …, *β*_*k*_. Given isotype transition probabilities **P**, the joint probabilities CSR(*T*_1_, *β*_1_, **P**), …, CSR(*T*_*k*_, *β*_*k*_, **P**) are conditionally independent, resulting in the joint likelihood

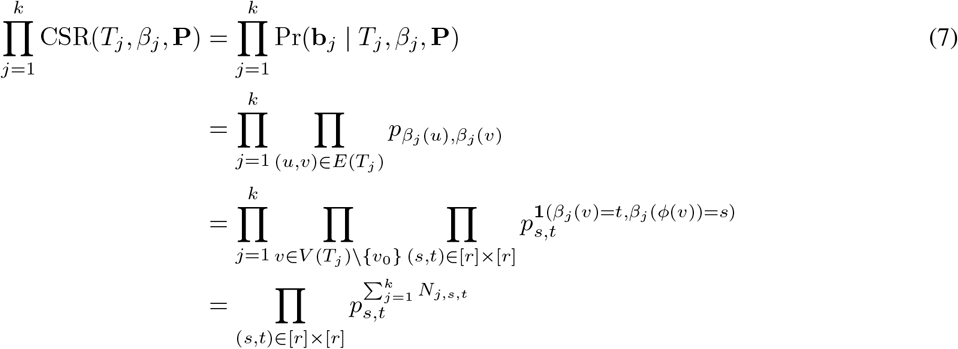

where *N*_*j,s,t*_ is the count of occurrences in lineage tree *T*_*j*_ such that *β*(*v*) = *t* and *β*(*ϕ*(*v*)) = *s*.

## B Combinatorial characterization and complexity results

### B.1 B cell lineage forest inference

Recall the B Cell Lineage Forest Inference Problem (BLFI) from Sec. 2, restated below for convenience.

**(Main Text) Problem 1** (B Cell Lineage Forest Inference (BLFI)). Given MSAs **A**_1_, …, **A**_*k*_ and isotypes **b**_1_, …, **b**_*k*_ for *k* clonotypes, find isotype transition probabilities **P**^*^ for *r* isotypes and lineage trees 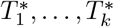for (**A**_1_, **b**_1_), …, (**A**_*k*_, **b**_*k*_) whose nodes are labeled by sequences 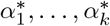 and isotypes 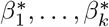, respectively, such that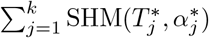 is minimum and then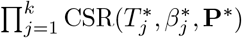 is maximum.

#### Theorem 1.

The BLFI problem is NP-hard even if *k* = 1 and *r* = 1.

We prove that the BLFI problem is NP-hard via a simple reduction from the Large Parsimony problem [41]. Although this problem is well known, we restate it here for completeness.

#### Problem 3

(Large Parsimony (LP)). Given a matrix **A** ∈ {0, 1}^*n×m*^, find a rooted tree *T* whose nodes are labeled by sequences *α* : *V* (*T*) → {0, 1}^*m*^ such that the *n* leaves are labeled by the rows of **A** and ∑_(*u,v*)*∈E*(*T*)_ *D*(*α*(*u*), *α*(*v*)) is minimium.

The reduction to BLFI proceeds by using the same MSA **A** directly for a single clonotype, i.e., *k* = 1. Additionally, we restrict the number *r* of isotypes to 1, and set isotypes **b** = [1]^*n*^.

#### Lemma 1.

Tree *T* and node labeling *α* form an optimal solution to LP instance **A** if and only if tree *T*, sequences *α* and isotypes *β*, the isotype transition probabilities **P** form an optimal solution to BLFI instance (**A, b**).

*Proof*. (⇒) Let tree *T* and sequence labeling *α* be an optimal solution to the LP problem. We will show that *T* and *α* can be augmented to form an optimal solution to the corresponding BLFI problem. We set **P** = [1]. We also set *β*(*v*) = 1 for all nodes *v ∈ T*. We claim that (*T, α, β*, **P**) form an optimal solution to BLFI. Assume for a contradiction there exists a solution (*T* ′, *α*′, *β*′, **P**′) such that SHM(*T*′, *α*′) < SHM(*T, α*), or SHM(*T*′, *α*′) = SHM(*T, α*) and CSR(*T*′, *β*′, **P**′) *>* CSR(*T, β*, **P**). Clearly, any feasible solution to BLFI must use *β*(*v*) = 1 for all nodes *v* and **P** = [1] as *r* = 1. This means that any feasible solution to BLFI will have a CSR objective value of 1. Therefore, CSR(*T*′, *β*′, **P**′) = CSR(*T, β*, **P**) = 1. Hence, SHM(*T*′, *α*′) < SHM(*T, α*). As can be seen in (1), the SHM objective equals the objective of the LP problem. Therefore, *T* ′ and *α*′ have a lower parsimony score than *T* and *α*, a contradiction.

(⇐) Let (*T, α, β*, **P**) be an optimal solution to BLFI. Again, as the SHM objective equals the objective of the LP problem, it directly follows that (*T, α*) form an optimal solution to the LP problem instance. □

### B.2 Most parsimonious tree refinement

#### B.2.1 Combinatorial characterization

Recall from the main text the definition of isotype transition probabilities **P**, the CSR log-likelihood for isotypes **b** of a tree *T* with nodes labeled by isotypes *β*, and the Most Parsimonious Tree Refinement problem, provided below for convenience.

**(Main Text) Definition 2**. An *r* × *r* matrix **P** = [*p*_*s,t*_] is an *isotype transition probability matrix* provided for all isotypes *s, t* ∈ [*r*] it holds that (i) *p*_*s,t*_ ≥ 0, (ii) *p*_*s,t*_ = 0 if *s > t*, and 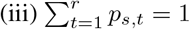 for all isotypes *s* ∈ [*r*].

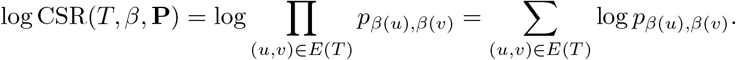

**(Main Text) Problem 2** (Most Parsimonious Tree Refinement (MPTR)). Given a tree *T* on *n* leaves, isotypes **b** = [*b*_0_, …, *b*_*n*_] and isotype transition probabilities **P**, find a tree *T* ′ with root 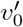 and isotype labels *β*′ : *V* (*T* ′) → [*r*] such that (i) *T* ′ is a refinement of *T*, (ii)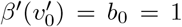, (iii)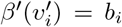 for each leaf 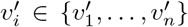 and (iv) log CSR(*T*′, *β*′, **P**) is maximum.

Let *σ* be a mapping from *V* (*T* ′) to *V* (*T*) that reverses all expand operations of each node *u*′ in refinement *T* ′ in order to obtain back the node *σ*(*u*′) = *u* from which it was derived in the original tree *T*. We say that an isotype labeling *β*′ : *V* (*T* ′) → [*r*] of *T* ′ is transitory if along each directed edge (*u*′, *v*′) of *T* ′ either the isotype changes or *u*′ and *v*′ correspond to two distinct nodes of *T*. More formally, we have the following definition.

##### Definition 3.

Let *T* ′ be a refinement of a tree *T* whose leaves are labeled by isotypes **b**. Then, an isotype labeling *β*′ of *T* ′ is *transitory* provided 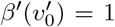 where 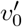 is the root of *T* ′, 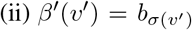 for each leaf *v*′*∈ L*(*T*′), (iii) *β*′(*u*′)≤ *β*(*v*′) for each edge (*u*′, *v*′) of *T* ′, and (iv) *β*′(*u*′) = *β*′(*v*′) only if *σ*(*u*′)≠*σ*(*v*′) for each edge (*u*′, *v*′) of *T* ′.

Importantly, among the set of optimal solutions (*T* ′, *β*′) to each MPTR problem instance (*T*, **b, P**) there exist solutions where *β*′ is transitory.

##### Lemma 2.

Let (*T*, **b, P**) be an MPTR problem instance. There exist an optimal solution (*T* ′, *β*′) where *β*′ is transitory.

*Proof*. We prove this by contradiction. Let (*T* ′, *β*′) be an optimal solution where *β*′ is not transitory. First, observe that it holds that *β*′(*u*′)≤ *β*(*v*′) for each edge (*u*′, *v*′) of *T* ′. To see why, if there were an edge (*u*′, *v*′) such that *β*′(*u*′) *> β*(*v*′) then CSR(*T*′, *β*′, **P**) = −∞ as log *p*_*s,t*_ = if *s > t*. However, setting *β*′(*u*′) = 1 for nodes *u*′ would result in log-likelihood greater than −∞. Since (*T* ′, *β*′) is a feasible solution to MPTR respecting irreversibility of isotype transitions, it means that condition (iv) of Definition 3 is violated. Let (*u*′, *v*′) be an edge such that *β*′(*u*′) = *β*′(*v*′) and *σ*(*u*′) = *σ*(*v*′). We can contract this edge, retaining the isotype labeling *β*′ for the remaining nodes, such that the resulting tree remains a refinement of *T* and the objective value remains unchanged as log *p*_*s,s*_ = 0. Repeating this procedure for all edges (*u*′, *v*′) such that *β*′(*u*′) = *β*′(*v*′) and *σ*(*u*′) = *σ*(*v*′) results in (*T* ″, *β*″), where *T* ″ is a refinement of *T* labeled by *β*″, with the same optimal score as (*T* ′, *β*′). Clearly, (*T* ″, *β*″) is transitory, proving the lemma.

#### B.2.2 Complexity

Note that maximizing the CSR log-likelihood is equivalent to maximizing the CSR likelihood, which is the objective function we will use in this subjection. That is,

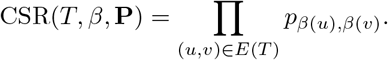

We now prove the following theorem.

##### Theorem 2.

The MPTR problem is NP-hard.

We show that MPTR is NP-hard by reduction from Set Cover.

##### Problem 4

(Set Cover). Given a universe𝒰of elements {*u*_1_, … *u*_|𝒰|_} and a collection 𝒮 of subsets {*S*_1_, …, *S*_|𝒮|_} such that 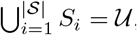, find a cover 𝒞 ⊆ 𝒮 such that *U* _𝒮*∈*𝒞_ *S* = 𝒰 and the size | 𝒞| of the cover is minimum.

Note that while the order of the subsets in collection does not matter for Set Cover, our reduction will assume the subsets to be in an arbitrary but fixed order. Similarly, we will 𝒰assume to be ordered arbitrarily. Set Cover has been proven to be NP-hard in Karp’s 21 NP-complete problems [42]. We describe a polynomial time reduction from Set Cover to MPTR. To that end, given the set 𝒰of elements and the collection of subsets, we construct a tree *T* with |𝒰 |+ 1 leaves, *r* = |𝒰 |+|𝒮| + 2 isotypes, observed isotypes **b***∈* [*r*]^|^ 𝒰^|+1^, and *r × r* transition probabilities **P**. The steps are as follows.

1. To construct tree *T*, we begin by adding the root node *v*_0_. Following that, we attach two children, denoted as 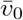 and *v*_|*𝒰*|+1_, to the root node *v*_0_. Finally, for each element *u*_*q*_ *∈* 𝒰, we add an edge (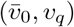 in tree *T*. The constructed tree *T* has | 𝒰| + 3 nodes and | 𝒰| + 2 edges.
2. We consider a total of *r* = | 𝒮| + | 𝒰| + 2 isotypes, each corresponding to either a subset *S*_*i*_ *∈* 𝒮, an element *u*_*q*_ *∈* 𝒰, or one of the special symbols ⊤ or ⊥. Specifically, the first isotype stands for the special symbol ⊤, followed by | 𝒰| isotypes representing each subset *S*_*i*_ *∈* 𝒮, succeeded by | 𝒰| isotypes representing each element *u*_*q*_ *∈* 𝒰, and concluding with the last isotype signifying the special symbol ⊥. For convenience, we define a function *R* : 𝒮 ∪ 𝒰 ∪ {⊤, ⊥} → [*r*] to map the subsets *S*_*i*_ *∈* 𝒮, the elements *u*_*q*_ *∈* 𝒰, and the special symbols ⊤ and ⊥ to their representative isotype indices as follows.

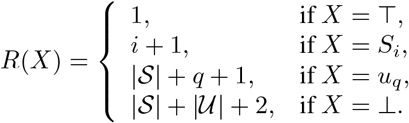
3. For the observed isotypes, we set *b*_0_ = *b*_| *𝒰*|+1_ = *R*(⊤) = 1, and *b*_*q*_ = *R*(*u*_*q*_) for 1 ≤ *q* ≤ | 𝒰|.
4. We define *ϵ* to be a constant such that 0 < *ϵ* ≤ 1*/*(|𝒮| +|𝒰| + 1). Next, we construct the isotype transition probabilities **P** parameterized by *ϵ* as follows.
  a. We set the transition probability from *R*(⊤) to *R*(⊤) or *R*(*S*_*i*_) for any set *S*_*i*_ *∈ S* to be *ϵ* and to *R*(*u*_*q*_) for any *u*_*q*_ *∈* 𝒰 to be 0.

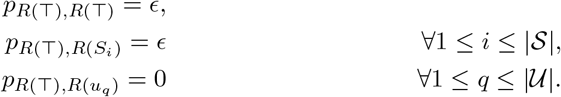
  b. We set the transition probability from *R*(⊤) to *R*(⊥) to be 1 − (1 + |𝒮|)*ϵ*.

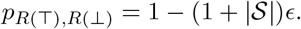
  c. We set the transition probability 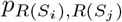 for any *S*_*i*_, *S*_*j*_ *∈* 𝒮 to be *ϵ* if *i* < *j*, and 0 otherwise.

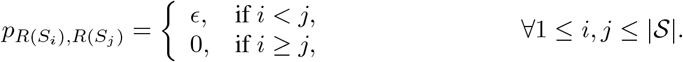
  d. We set the transition probability from *R*(*S*_*i*_) to *R*(*u*_*q*_) for any set *S*_*i*_ *∈* 𝒮 and any element *u*_*q*_ *∈* 𝒰 to be *ϵ* if *u*_*q*_ *∈ S*_*i*_, and 0 otherwise.

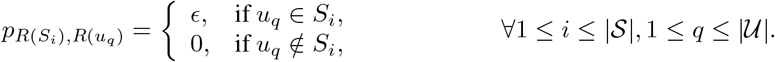
  e. For each *S*_*i*_ *∈ 𝒮*, we set the transition probability from *R*(*S*_*i*_) to *R*(⊤) to be 0 and to *R*(⊥) to be 1 − (|𝒮| − *i* + |*S*_*i*_|)*ϵ*.

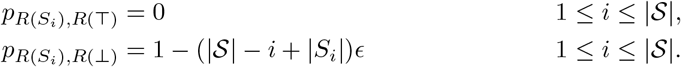
  f. For any *u*_*q*_ *∈* 𝒰, we set the transition probability from *R*(*u*_*q*_) to any other isotype except ⊥ to be 0. We set 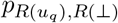 for any *u*_*q*_ *∈* 𝒰 to be 1.

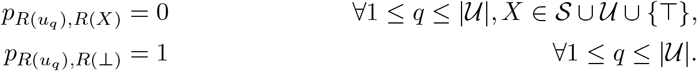
  g. Last, we set the transition probability *p*_*R*(⊥),*R*(⊥)_ to be 1.

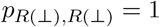

**Figure S1:**
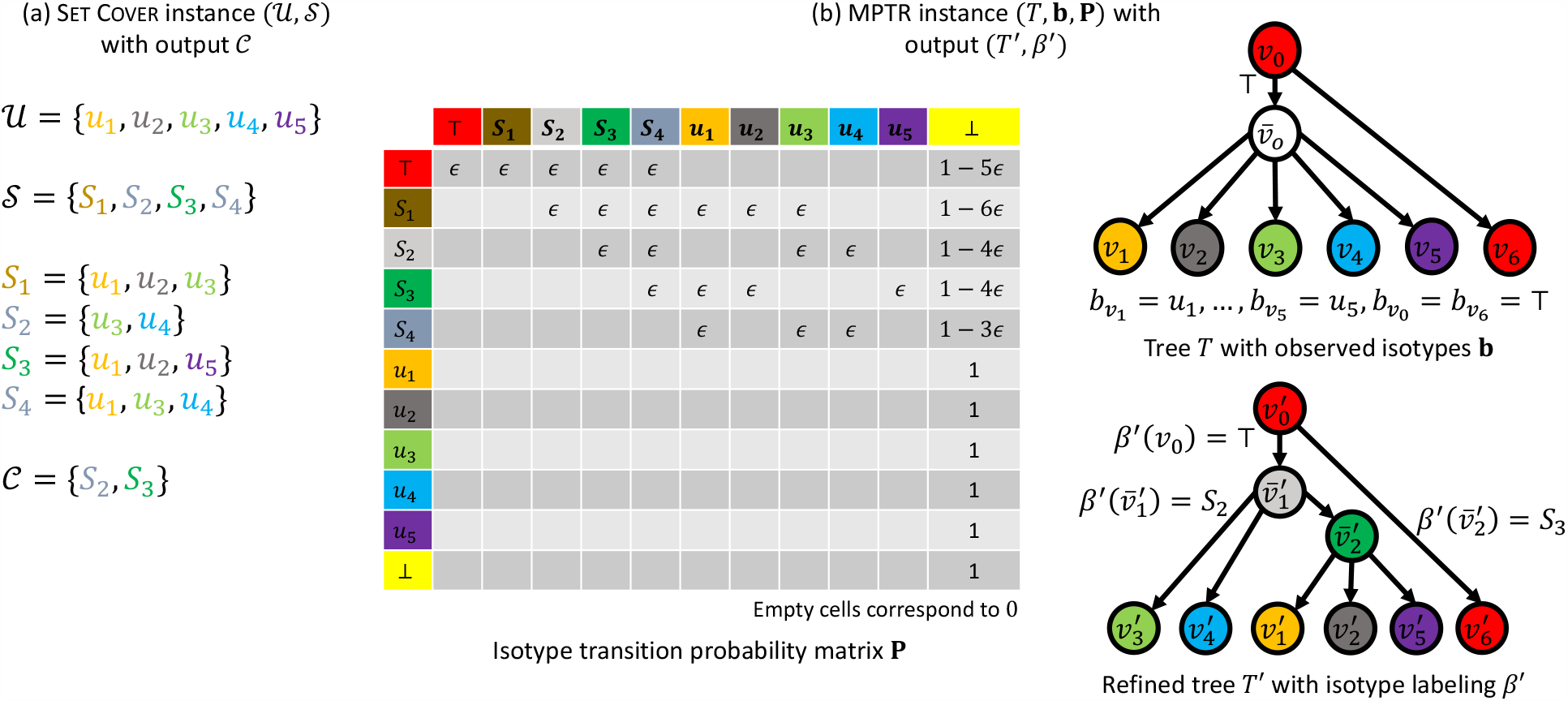
Polynomial time reduction from Set Cover to MPTR. (a) shows a Set Cover instance (𝒰, 𝒮), with the corresponding minimum set cover 𝒞. The constructed MPTR instance (*T*, **b, P**), along with the output (*T* ′, *β*′) is shown in (b). Isotypes are indicated through colors. The mapping function *R* is omitted, with the isotypes directly represented by elements, subsets, *⊤*, or *⊥*. The empty boxes in the transition probability matrix **P** corresponds to 0.

Clearly, by construction matrix **P** obtained from a Set Cover instance (𝒰, 𝒮) is an isotype transition probability matrix as **P** is upper triangular, each entry is non-negative and each row sums to 1. In addition, this reduction takes polynomial time.

To prove hardness, let (*T* ′, *β*′) be an optimal solution to the MPTR instance composed of the input tree *T*, observed isotypes **b**, and isotype transition probabilities **P** corresponding to Set Cover instance (𝒰, 𝒮).

##### Lemma 3.

CSR(*T*′, *β*′, **P**) *>* 0 for the refined tree *T* ′ and the isotype labeling *β*′ inferred by MPTR.

*Proof*. We prove this by showing that for any constructed input tree *T*, observed isotypes **b** and isotype transition probabilities **P**, there exists a refined tree *T* ′ and isotype labeling *β*′ such that CSR(*T*′, *β*′, **P**) *>* 0. We provide a proof by constructing a refined tree *T* ′ with isotype labeling *β*′. The tree *T* ′ will expand the unique polytomous node 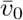 into a chain 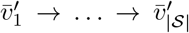. We leave the remaining nodes *v*_0_, *v*_1_, …, *v*_|*𝒰*|+1_ of *T* unaltered, letting 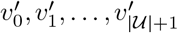 that denote their corresponding nodes in *T* ′. Next, for each 1 ≤ *q* ≤ | 𝒰|, we pick a subset *S*_*i*_ such *u*_*q*_ *∈ S*_*i*_, and add edge 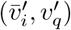 in *T* ′ and set 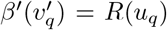. We add the edges 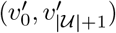 and 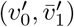. Finally, we set 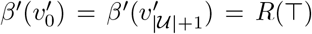. Clearly all the edges in *T* ′ have nonzero isotype transition on probabilities, so CSR(*T*′, *β*′,**P**) > 0. □

##### Corollary 2.

The root 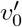 of *T* ′ is labeled by isotype ⊤.

*Proof*. Due to the presence of leaf *v*_|*𝒰*|+1_ with isotype *b*_|*𝒰*|+1_ = *R*(*⊤*), the root 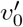 of *T* ′ must be labeled by isotype 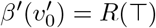, otherwise there would be a zero-probability edge.

##### Corollary 3.

No node *v*′ of *T* ′ is labeled by isotype ⊥.

##### Corollary 4.

Each edge (*v*′, *v*″) of *T* ′ has an isotype transition probability of *p*_*β*′(*v*′),*β*′(*v*″)_ = *ϵ*.

Observe that 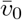 is the only polytomous node in *T*. We will now prove that 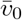 is the only node of *T* that is expanded in the refined tree *T* ′.

##### Lemma 4.

Node 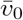 is the only node of *T* that is expanded in *T* ′.

*Proof*. By Lemma 2, we may assume that *β*′ is transitory. Let 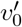 be the root of *T* ′. We prove this lemma by contradiction. Let 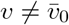 be a distinct node of *T* that is expanded in *T* ′. We distinguish the following three cases.

- *v* = *v*_|*𝒰*|+1_: In this case, *v* equals the leaf node *v*_|*𝒰*|+1_ whose parent is the root *v*_0_. Consider the corresponding node 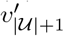 of *T*′ such that 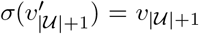 and 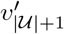 is a leaf of *T*′. Since *β*′ is transitory, we have that 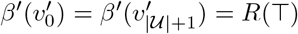. Since node *v*_|*𝒰*|+1_ was expanded, node 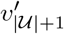 has a unique parent 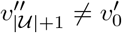. As *β*′ is transitory and 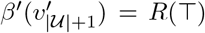 and *R*(*⊤*) *≤ s* for all *s* ∈ [*r*], we must have that 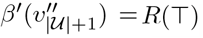. This, however, implies that *β*′ is not transitory as 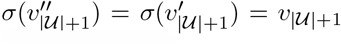 and 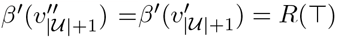, which yields a contradiction.
- *v* ∈{*v*_1_, …, *v*_|*𝒰*|_}: Note that *v* is a leaf of *T*. Consider the corresponding node *v*′ of *T* ′ such that *σ*(*v*′) = *v* and *v*′ is a leaf of *T* ′. The parent of *v* in *T* is node 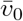. Since node *v* was expanded, node *v*′ has a unique parent *v*″ such that *σ*(*v*″) = *v*. Let *v*^*″′*^ be the unique parent of *v*″. By Corollary 4, we have that the two edges (*v*″, *v*′) and (*v*^*‴*^, *v*″) both have probabilities *ϵ*, contributing a factor of 2*ϵ* to the overall probability CSR(*T*′, *β*′, **P**). However, by contracting the edge (*v*″, *v*′) and removing the node *v*″, we obtain another solution with higher probability, leading to a contradiction.
- *v* = *v*_0_: Consider the corresponding node 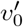 such that 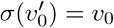 and 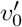 is the root of *T* ′. There are two cases two consider. Let 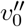 be a child of 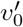 such that 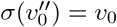. We distinguish two cases.
  - First, 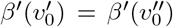. By Corollary 2, we have that 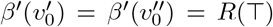. By Corollary 4, we have that the edge 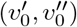 contributes a factor of *ϵ* to the overall probability CSR(*T*′, *β*′, **P**). We can remove this factor by simply contracting the edge 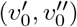, resulting in a more optimal solution, which is a contradiction.
  - Second, 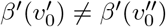. By Corollary 2, we have that 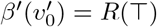. By Lemma 3, we have 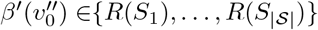. Again, by the same lemma, all children of 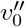 will be labeled by isotypes different than 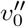. In particular, each child of 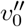 will either correspond to node *v*_0_ or 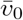 of *T*, labeled from the set 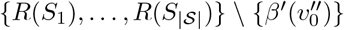. Thus, we may contract the edge 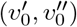, with probability *ϵ*, and remove the node 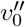, reassigning all children of 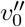 to 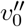. The resulting tree and isotype labeling will have a larger probability, a contradiction.

Assume that a series of expand operations on 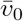 in *T* has generated *k* nodes in *T* ′, where *k* ranges from 1 (no expand operation) to |*𝒰*|. We denote 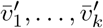 to be the new nodes in *T* ′ originating from 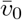 in *T*, i.e., 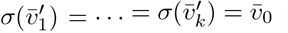. Let 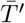 be the subtree of *T* ′ induced by nodes 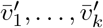.

##### Lemma 5.

The refined tree *T* ′ has |*𝒰*| + *k* + 2 nodes, |*𝒰*| + *k* + 1 edges, and CSR(*T*′, *β*′, **P**) = *ϵ*^|*𝒰*|+*k*+1^.

*Proof*. Since *T* has |*𝒰*| + 3 nodes, and, by Lemma 4, the only node 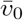 of *T* that is expanded, expands to *k* nodes 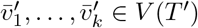, the total number of nodes in *T* ′ is |*𝒰*| + 2 *−* 1 + *k* = |*𝒰*| + *k* + 2. Similarly, the number of edges in *T* is |*𝒰*| + 2, and since 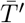 is a tree containing *k* nodes, it has *k −* 1 edges. So the total number of edges in *T* ′ is |*𝒰*| + 2 + *k −* 1 = |*𝒰*| + *k* + 1. It follows from Corollary 4 that CSR(*T*′, *β*′, **P**) = *ϵ*^|*𝒰*|+*k*+1^.

##### Lemma 6.

Nodes 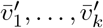 of *T* ′ are labeled by *k* distinct isotypes from the set {*R*(*S*_1_), …, *R*(*S*_| *𝒮* |_)}.

*Proof*. By construction of **P**, *R*(*u*_*q*_) can only be transitioned into from *R*(*S*_*i*_) with nonzero probability where *u*_*q*_ *∈ S*_*i*_. So if there is an edge 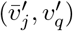 in *T* ′ connecting expanded node 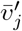 with leaf 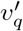 labeled with 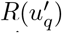 then 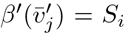 for some *S*_*i*_ *∈ 𝒮*. Using the observation, we begin by showing that each expanded node 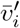 has at least one child 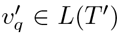. We do so by contradiction. Suppose the refined tree *T* ′ has an expanded node 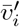 that does not have any leaf 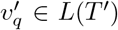 as a child. Without loss of generality, assume that 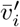 has a child 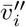, which, in turn, is the parent of a leaf 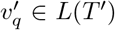. This means that 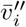 is labeled with 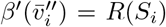 for some *S*_*i*_ *∈ S*. Since *R*(*S*_*i*_) can only be transitioned into from *R*(*S*_*j*_), where *j* < *i*, or *R*(⊤) with nonzero probability, it holds that 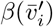 is either *R*(*S*_*j*_) where *j* < *i* or *R*(⊤). Similarly, the parent of 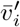 should also be labeled either with *R*(*S*_*j*_′) where *j*′ < *j* or *R*(⊤). Now we create a new tree *T* ″ by (i) adding the children of 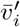 as the children of the parent of 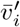, and (ii) deleting the edge between 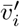 and its parent. Clearly *T* ″ has nonzero transition probabilities on all the edges, but has one fewer edge than *T* ′. So CSR(*T*″, *β*′, **P**) < CSR(*T*′, *β*′, **P**), which contradicts with the premise that *T* ′ minimizes CSR(*T*′, *β*′, **P**). So each expanded node 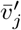 is labeled with *R*(*S*_*i*_) for some *S*_*i*_ *∈ 𝒮*.

It remains to show that the *k* nodes 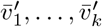 are labeled by *k* distinct isotypes from the set {*R*(*S*_1_), …, *R*(*S*_| *𝒮*|_)}. To see why, observe that, by construction of **P**, the incident nodes of each edge among nodes 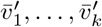 must be labeled by distinct isotypes from the set {*R*(*S*_1_), …, *R*(*S*_| *𝒮* |_)}, as *p*_*R*(*S*_^*i*^),*R*(*S*^*i*^) = 0 for all *S*_*i*_ *∈ 𝒮*.

##### Lemma 7.

There exists an minimum set cover of size *k* if and only if there is an optimal solution (*T* ′, *β*′) such that CSR(*T*′, *β*′, **P**) = *ϵ*^|*𝒰*|+*k*+1^.

*Proof*. (*⇒*) Let 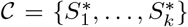 be a set cover of minimum size *k*. Without loss of generality, we further assume that 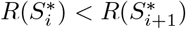 for any 1 *≤ i ≤ k −* 1. Next, we build a refined tree *T* ′ with isotype labeling *β*′ by expanding the node 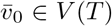 to *k* nodes 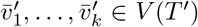. More specifically, we replace 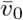 with 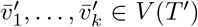 such that (i) *v*_0_ is connected to 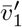 by an edge, (ii) there is an edge 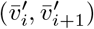 in *T* ′ for each 1 *≤ i ≤ k −* 1, (iii) 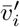 is labeled with 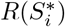, i.e. 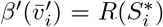, and (iv) for each child *v*_*q*_ of 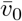 in *T*, there exists exactly one edge 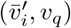 in *T* ′ where 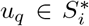. Clearly *T* ′ is a refinement of tree *T*, and all the newly added edges have nonzero transition probabilities *ϵ*. Hence, CSR(*T*′, *β*′, **P**) = *ϵ*^|*𝒰*|+*k*+1^.

All that remains to show is that (*T* ′, *β*′) is optimal. We show this by contradiction. Let (*T* ″, *β*″) be an optimal solution such that CSR(*T*″, *β*″, **P**) < CSR(*T*′, *β*′, **P**) = *ϵ*^|*𝒰*|+*k*+1^. By Lemma 4, we have that only the node 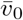 of *T* is expanded in *T* ″ corresponding 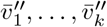′ nodes in *T* ″. Since CSR(*T*″, *β*″, **P**) < CSR(*T*′, *β*′, **P**), it must hold that *k*″ < *k*. By Lemma 6 we have that the *k*′ labels of nodes 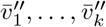′ correspond to *k*′ distinct subsets of 𝒮. By Lemma 3, we have that these *k*′ subsets of 𝒮 form a cover of the universe 𝒰, leading to a contradiction. Hence, (*T* ′, *β*′) is optimal.

(⇐) Now assume that there exists an optimal solution (*T* ′, *β*′) such that CSR(*T*′, *β*′, **P**) = *ϵ*^|*𝒰*|+*k*+1^. Note that the restriction that CSR(*T*′, *β*′, **P**) = *ϵ*^|*𝒰*|+*k*+1^ is without loss of generality due to Lemma 5. Now according to Lemma 6, there are *k* expanded nodes in *T* ′ labeled with 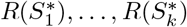. We define 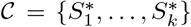. Now each leaf 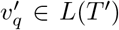 labeled with *R*(*u*_*q*_) is the child of an expanded node 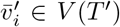 labeled with 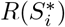. Since CSR(*T*′, *β*′, **P**) *>* 0 by Lemma 3, the transition probability from 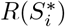 to *R*(*u*_*q*_) is strictly greater than 0, which means 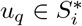. So every element in *𝒰* is covered by one of the subsets from 𝒞. So 𝒞 is a set cover of size *k*.

It remains to show that 𝒞 is a minimum-size set cover. Assume for a contradiction that there exists a cover 𝒞*′* =*⊆* 𝒮 such that |𝒞*′*| = *k*′ < *k* = |𝒞|. Let 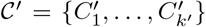 where the subsets follow the same order as in the original reduction to MPTR. We construct a refined tree *T* ″ with isotype labeling *β*″ corresponding to 𝒞*′* by expanding the unique polytomous node 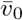 of *T* into a chain 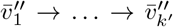, with one node 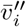 for each subset *C*_*i*_ *∈* 𝒞*′* labeled by 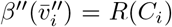, and connecting each leaf *v*_*q*_ *∈ {v*_1_, …, *v*_|*𝒰*|_} to a single expanded node 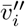 such that 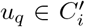. Since 𝒞′ is a cover of *𝒰*, each leaf *v*_*q*_ ∈ {*v*_1_, …, *v*_|*𝒰*|_} will be connected. Moreover, tree *T* ″ with isotype labeling *β*″ form a solution to MPTR. Clearly, *T* ″ has + *k*′ + 2 nodes and + *k*′ + 1 edges. Moreover, each edge of *T* ″ has a nonzero isotype transition probability equal to *ϵ*, so CSR(*T*″, *β*″, **P**) = *ϵ*^|*𝒰*|+*k*^*′*+1 < *ϵ*^|*𝒰*|+*k*+1^ = CSR(*T*′, *β*′, **P**), a contradiction.

## C Supplementary Methods

**Figure S2:**
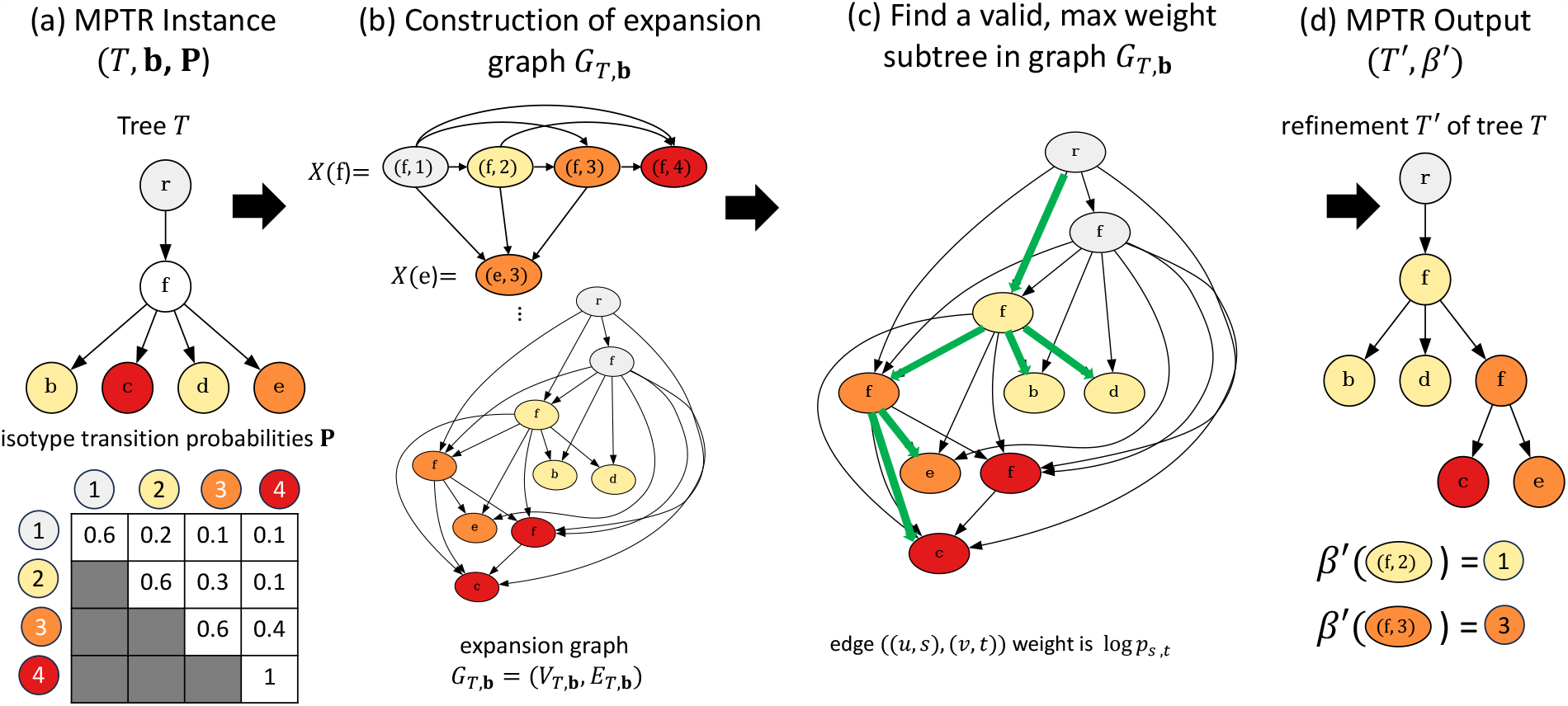
Algorithm for solving the MPTR problem. (a) An instance (*T*, **b, P**) of the MPTR problem. (b) To construct the expansion graph *G*_*T*,**b**_ for tree *T* whose leaves have isotypes **b**, each original node *u* in *V* (*T*) corresponds to a set *X*(*u*) of nodes in *G*_*T*,**b**_. Edges are added to capture all transitory refinements of tree *T*. (c) We use the expansion graph *G*_*T*,**b**_ with weighted edges to find a valid, maximum weight subtree in *G*_*T*,**b**_, depicted in green. (d) This selected subtree is an optimal solution (*T* ′,*β*′) to the MPTR problem instance (*T*, **b, P**).

### C.1 Tree refinement

We solve an instance (*T*, **b, P**) of the MPTR problem (Fig. S2a) by reducing it to a graph problem. Given an instance (*T*, **b, P**) of the MPTR, we construct a directed graph *G*_*T*,**b**_, called the expansion graph, with nodes *V* (*G*_*T*,**b**_) ⊆ *V* (*T*) *×* [*r*] and edges *E*(*G*_*T*,**b**_). At a high level, nodes of *V* (*G*_*T*,**b**_) are of the form (*u, s*) where *u ∈ V* (*T*) is a node of the input tree *T* and *s* ∈ [*r*] is an isotype state. Formally, we have the following definition.

#### Definition 4.

A directed graph *G*_*T*,**b**_ is an *expansion graph* of a rooted tree *T* whose leaves are labeled by isotypes **b** provided *V* (*G*_*T*,**b**_) =⋃*u∈V* (*T*) *X*(*u*) where

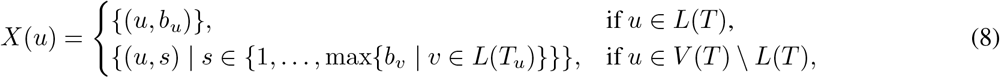

and *E*(*G*_*T*,**b**_) = {((*u, s*), (*v, t*)) | (*u, v*) *∈ E*(*T*), *s ≤ t} ∪ {*((*u, s*), (*u, t*)) | *u ∈ V* (*T*), *s* < *t}*.

In the above definition *X*(*u*) is the set of nodes of *G*_*T,b*_ corresponding to node *u* of *T*, accounting for the fact that leaves *u* of *T* retain their isotype state in any refinement *T* ′ of *T*. On the other hand, internal nodes *u* of *T* may be subject to expand operations such that the corresponding nodes of *T* ′ are assigned isotypes *s* ranging from state 1 to the maximum isotype state among all descendant leaves of *u* in *T*. The edges of *G*_*Tb*,_ respect the irreversibility property of isotypes as well as the parental relationships of nodes of *T*. See Fig. S2c for an example expansion graph *GT,b*.

We now define constrained subtrees, termed valid, of the expansion graph *G*_*T,b*_.

#### Definition 5.

A subtree *T* ′ of *G*_*T,b*_ is *valid* provided (i) *T* ′ is rooted at (*v*_0_, 1) where *v*_0_ is the root of *T* and (ii) there is a unique edge ((*u, s*), (*v, t*)) in *E*(*T*′) for each edge (*u, v*) of *T*.

We now show that the set of valid subtrees of *G*_*T*,**b**_ corresponding to trees *T* ′ with isotype labelings *β*′ is equivalent to the set composed of pairs (*T* ′, *β*′) where *T* ′ is a refinement of *T* and *β*′ is a transitory isotype labeling of *T* ′.

#### Lemma 8.

Let *T* ′ be a refinement of *T* whose leaves are labeled by isotypes **b** and let *β*′ be an isotype labeling of *T* ′. Then, *β*′ is transitory if and only if (*T* ′, *β*′) induces a valid subtree of *G*_*T,b*_.

*Proof*. (*⇒*) Let *β*′ be a transitory isotype labeling of *T* ′. We start by showing that (*T* ′, *β*′) induce a connected subtree of *G*_*T*,**b**_. First, let *u*′ be a node of *T* ′ labeled by isotype *β*(*u*′). We claim that (*u*′, *β*(*u*′)) *∈ X*(*u*). We distinguish the two cases. First, *u*′ *∈ L*(*T*′). Let *u* = *σ*(*u*′) be the original leaf node *u* of *T*. Since *β*′ is transitory, we have *β*(*u*′) = *b*_*σ*(*u*_*′*) = *b*_*u*_. Hence, (*u*′, *β*(*u*′)) *∈ X*(*u*) for each leaf node *u*′ *∈ L*(*T*′). Second, *u*′ *∈ V* (*T* ′) *\ L*(*T*′). Let *u* = *σ*(*u*′) be the original internal node *u* of *T*. Suppose for a contradiction (*u*′, *β*′(*u*′)) ∉ *X*(*u*)). This means that *β*′(*u*′) *>* max{*b*_*v*_ *∈ L*(*T*_*u*_)}. As such, there would be an edge (*u*″, *v*″) such that *β*′(*u*″) *> β*′(*v*″) where *u*″ is a node in the subtree 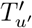 rooted at node *u*′. However, this would mean that *β*′ would violate condition (iii) of Definition 3, a contradiction. Thus, (*u*′, *β*(*u*′)) ∈ *X*(*u*) for each internal node *u*′ *V* (*T* ′\ *L*(*T*′). Hence, (*u*′, *β*(*u*′)) ∈ *V* (*G*_*T,b*_).

We now prove that each edge (*u*′, *v*′) of *T* ′ whose incident nodes are labeled by (*β*′(*u*′), *β*′(*v*′)) corresponds to an edge ((*u*′, *β*′(*u*′)), (*v*′, *β*′(*v*′))) of *G*_*T,b*_. This follows directly from conditions (iii) and (iv) of Definition 3 and the definition of *E*(*G*_*T,b*_) in Definition 4. This implies that the subgraph of *G*_*T,b*_ induced by (*T* ′, *β*′) is a (connected) subtree of *G*_*T,b*_.

We now must show that this induced subtree of *G*_*T,b*_ is valid. By condition (i) of Definition 3, we have that 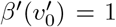 for the root 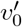 of *T* ′. As such, the induced subtree of *G*_*T,b*_ is rooted at 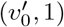. Finally, we must show there is a unique edge ((*u, s*), (*v, t*)) in the induced subtree of *G*_*T,b*_ for each original edge (*u, v*) of *T*. This follows from the fact that *T* ′ is a refinement of *T*. Thus the subgraph of *G*_*T,b*_ induced by (*T* ′, *β*′) is a valid subtree of *G*_*T,b*_.

(⇐) Consider a valid subtree of *G*_*T,b*_, resulting in a tree *T* ′ and isotype labeling *β*′. To see why *T* ′ is a refinement of *T*, observe that edges ((*u, s*), (*u, t*)) correspond to an expand operation on node *u* of *T*. It remains to show that *β*′ is transitory. By condition (i) of Definition 5, we have that the root of *T* ′ is labeled by state 1, satisfying condition (i) of Definition 3. Conditions (ii) and (iii) of Definition 3 are met by construction of *G*_*T*,**b**_. Finally, condition (iv) of Definition 3 follows from condition (ii) of Definition 5. Hence, the isotype labeling *β*′ of *T* ′ is transitory. The following key proposition follows from the previous two lemmas.

#### Proposition 2.

Let *G*_*T*,b_ be an expansion graph of a rooted tree *T* whose leaves are labeled by isotypes **b**. Then, given isotype transition probablities **P**, a valid subtree (*T* ′, *β*′) of *G*_*T,b*_ maximizing∑(*u*′,*v*′)*∈E*(*T*′) log *p*_*β*_′(*u*′),*β*′(*v*′) is an optimal solution to MPTR instance (*T*, **b, P**).

To find such a valid subtree with maximum log-likelihood, we formulate the following MILP based on a multi-commodity flow formulation for modeling connectivity. We make use of two sets of decision variables. The first is 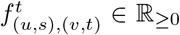, which represents the amount of flow on edge (*u, v*) designated for sink *q ∈ L*(*T*). The second is *x*_(*u,s*),(*v,t*)_ *∈ {*0, 1}, which indicates if edge (*u, v*) has non-zero flow.

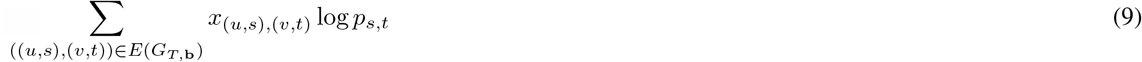

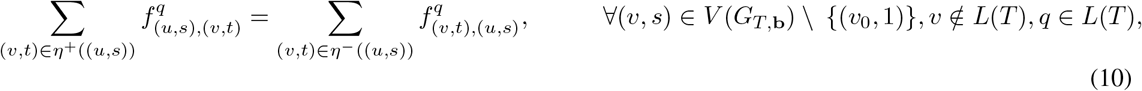

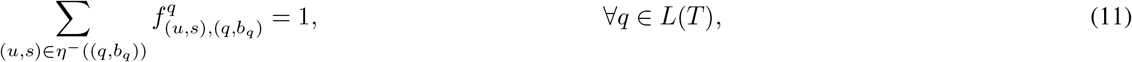

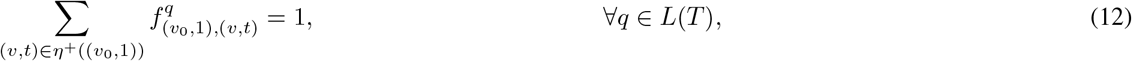

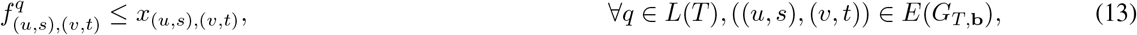

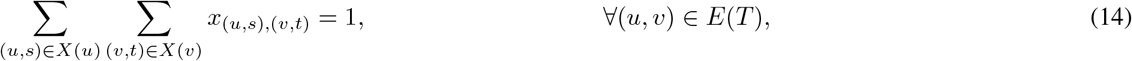

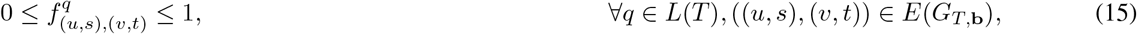

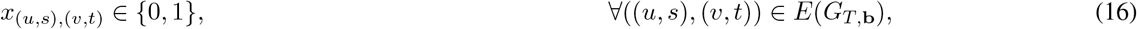

where *η*^+^((*u, s*)) is the set of direct successors of node (*u, s*) in graph *E*(*G*_*T*,**b**_) and *η*^*−*^((*u, s*)) is the set of direct predecessors of node (*u, s*).

Constraints (10), (11), (12) enforce flow conversation and ensure that each terminal receives one unit of flow. Below is a description of each of the above constraints. Constraint (13) links the flow variables to the choice of edges in the resulting refinement. Finally, constraint (14) ensures that refined tree *T* ′ can be obtained from tree *T* via a series of expand operations.

### C.2 Maximum likelihood estimate of isotype transition probabilities

Given a forest *T*_1_, …, *T*_*k*_ of lineage trees correspondingly labeled by isotypes *β*_1_, …, *β*_*k*_, we seek the maximum likelihood estimate of isotype transition probabilities **P**^*\**^.

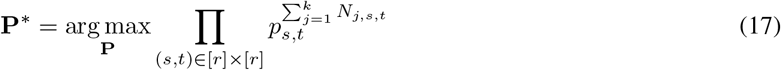

subject to

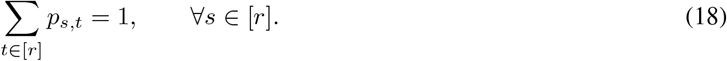

We solve this constrained optimization problem using Lagrange multipliers *λ*_*s*_ for each state *s*. We first take the log of likelihood 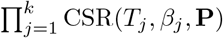 with respect to isotype transition probabilities **P**.

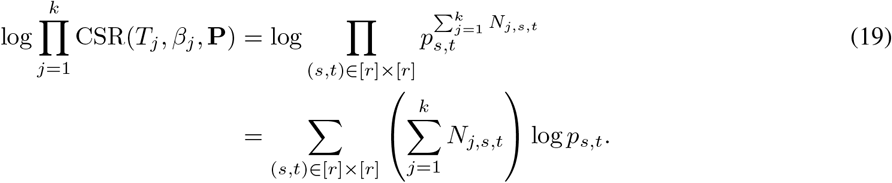

To our log-likelihood, we add the term *λ*_*s*_ ∑_*s∈*[*r*]_ *p*_*s,t*_ *−* 1 for each isotype *s*, resulting in new objective

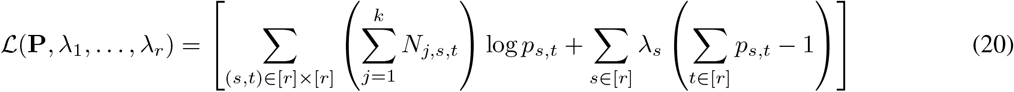

Then, we set the partial derivative of ℒ(**P**, *λ*_1_, …, *λ*_*r*_) with respect to each parameter *p*_*s,t*_ and *λ*_*s*_ and solve the resulting system of equations. For each *λ*_*s*_, we obtain our constraint,

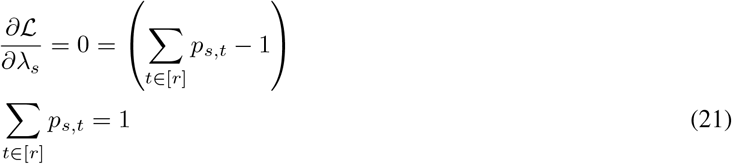

For each parameter *p*_*s,t*_, we set the partial derivative to 0 and solve for *p*_*s,t*_ as a function of *λ*_*s*_.

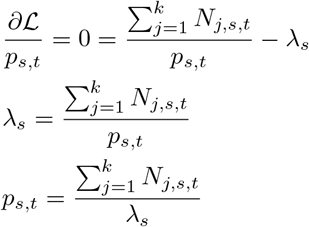

Given the constraint (21), we have that

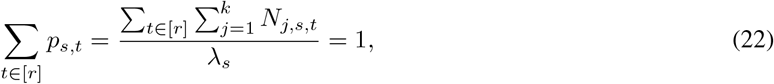

and

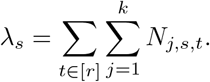

This yields the following maximum likelihood estimate 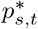,

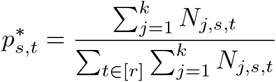

Lastly, to account for unobserved isotype transitions where isotype *s*≤*t*, we add a pseudocount of 1, resulting in updated isotype transition probabilities

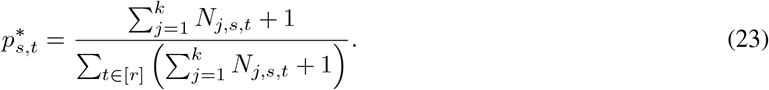

In the main text, we additionally use the shorthand 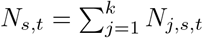.

## D Simulation details

We designed *in silico* experiments to evaluate TRIBAL with known ground-truth isotype transition probabilities **P** and lineage trees *T* labeled by sequences *α* and isotypes *β*. Specifically, we used an existing BCR phylogenetic simulator [13] that models SHM)(Appendix D.1 but not CSR. We generated isotype transition probabilities **P** with *r* = 7 isotypes (as in mice) under two different models of CSR (Appendix D.2). Briefly, both CSR models assume the probability of not transitioning is higher than the probability of transitioning, but in the *sequential model* there is clear preference for transitions to the next contiguous isotype, while in the *direct model* the probabilities of contiguous and non-contiguous class are similar (Fig. S3) Given **P**, we evolved isotype characters down each ground truth lineage tree *T*.

We generated 5 replications of each CSR model for *k* = 75 clonotypes and *n* ∈{ 35, 65} cells per clonotype, resulting in 20 *in silico* experiments, yielding a total of 1500 ground truth lineage trees. We generated our *in silico* experiments to evaluate all aspects of TRIBAL while benchmarking against existing methods including dnapars [7], dnaml [7] and IgPhyML [10].

### D.1 SHM simulation and benchmarking

The Davidsen and Matsen SHM simulator models the generation of B cell lineage trees via a Poisson branching process with selection towards BCRs with increased affinity [13]. We used the provided Docker Hub image container ^1^ to generate our ground truth B cell lineage trees *T* and sequence labels *α*. In addition, we used the provided benchmarking pipeline to run dnapars [7], dnaml [7] and IgPhyML [10]. Below is the command to generate our *in silico* experiments for *n ∈ {*35, 65} cells and *k* = 75 clonotypes and run comparison methods.

~~~
    simulate
        --igphyml
        --dnapars
        --dnaml
        --selection
        --target_dist=5
        --target_count=100
        --carry_cap=1000
        --T=35
        --lambda=2.0
        --lambda0=0.365
        --n={n}
        --nsim={k}
        --random_naive=sequence_data/AbPair_naive_seqs.fa
~~~

### D.2 CSR simulation

After generating each ground truth B cell lineage tree *T* as described above, we then evolved isotype characters down each tree *T* using two different models for class switch recombination to obtain ground truth isotypes *β*. First, we describe the two different CSR models that we used to generate ground truth isotype transition probabilities **P**. Then, we describe the generation of these isotype transition probability matrices under these two models.

We grouped each isotype transition probability *p*_*s,t*_ where *s* <= *t* into one of three categories: (i) *stay*, (ii) *next*, and (iii) *jump*(Fig S3a). In *stay*, the B cell does not undergo any class switching and the isotype does not change. In *next*, a B cell class switches to the next contiguous heavy chain locus. In *jump*, the B cell class switches by jumping to an isotype heavy chain constant locus that is not contiguous.

Next, we describe how we generated ground truth isotype transition probabilities **P** under both direct and sequential CSR models. To simulate isotype transition probabilities with direct switching, we randomly sampled a probability of transitioning 1 *− θ ∈ {*0.1, 0.15, …, 0.35}. We then set the initial isotype transition probabilities as

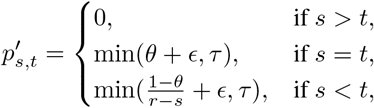

where we add Gaussian noise *ϵ∼𝒩* (*µ, σ*) with mean *µ* = 0.05 and standard deviation *σ* = 0.025 to each parameter. To avoid negative transition probabilities we set *τ* = 0.01. Fig. S3b shows an example of a simulated isotype transition probability matrix under the direct CSR model.

**Figure S3:**
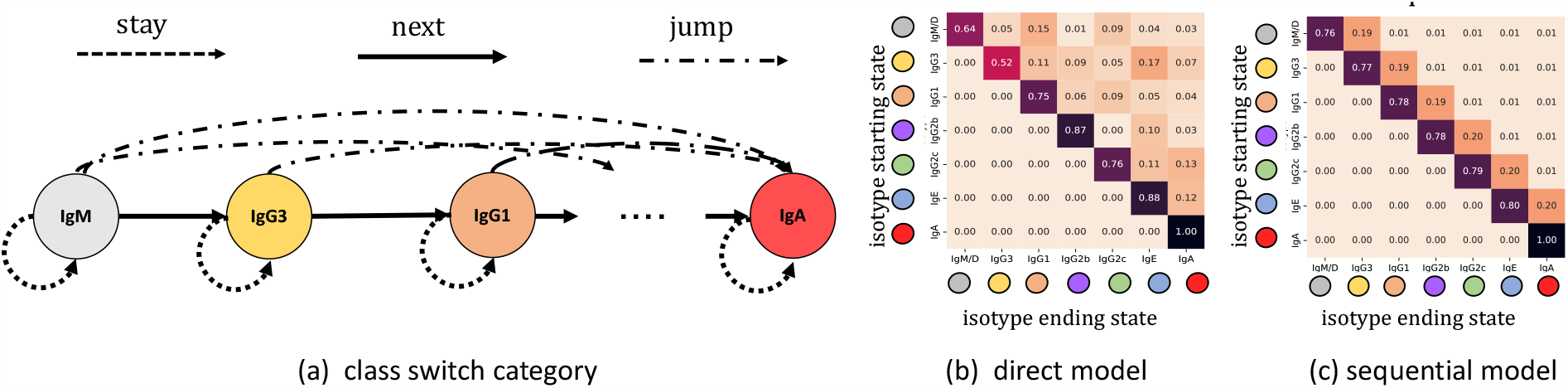
Class switch recombination models for *in silico* experiments. a) Examples of different isotype transition probability parameter groups. (b) Examples of simulated isotype transition probabilities **P** for the direct model of CSR. In the direct model, when a B cell class switches is no systematic preference for transition to the *next* sequential state or *jumping* to a non-contiguous isotype. (c) In the sequential model, a B cell undergoing CSR has a strong affinity for the *next* contiguous heavy chain locus.

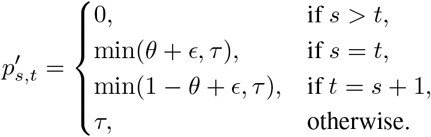

We then set parameter 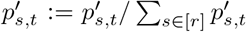 to ensure each row in the isotype transition probability matrix **P** sums to 1. Fig. S3b shows an example of a simulated isotype transition probability matrix under the direct model. Fig. S3c shows an example of a simulated isotype transition probability matrix under the sequential CSR model.

### D.3 Inference using TRIBAL

We ran TRIBAL in two ways, referred to as TRIBAL and TRIBAL-NO Refinement, in order to assess the importance of the tree refinement stage of our algorithm. As the naming convention implies, the main difference between TRIBAL and TRIBAL-NO Refinement, is that in TRIBAL-NO Refinement the input trees are not refined and the isotypes 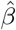 are inferred using the Sankoff [29] algorithm with weights 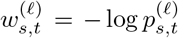. All other steps of TRIBAL algorithm remain the same.

Due to large input sets 𝒯_*j*_ for some simulated clonotypes *j*, we sample 50 trees from𝒯_*j*_ for consideration of candidate lineage tree 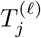 within each iteration *ℓ*. We additionally include the previous optimal lineage tree *T* ^(*ℓ−*1)^ of iteration *ℓ −*1 in the sampled trees for each clonotype *j* to ensure convergence.

We used a convergence threshold of 0.5 and a maximum of 10 iterations per restart. A total of 5 restarts were performed by iterating through *θ ∈ {*0.55, 0.65, 0.75, 0.85, 0.95} for each restart.

### D.4 Performance metrics

To evaluate performance of isotype transition probability inference, 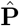, we utilized *Kullback-Leibler (KL) divergence*. To assess accuracy of lineage tree inference, we used normalized Robinson-Foulds (RF) distance to assess accuracy of the topology of the inferred tree 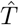, most recent common ancestor (MRCA) distance to assess accuracy of the inferred sequences 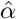, and Class Switch Recombination (CSR) error to assess accuracy of the inferred isotypes 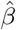.

**Figure S4:**
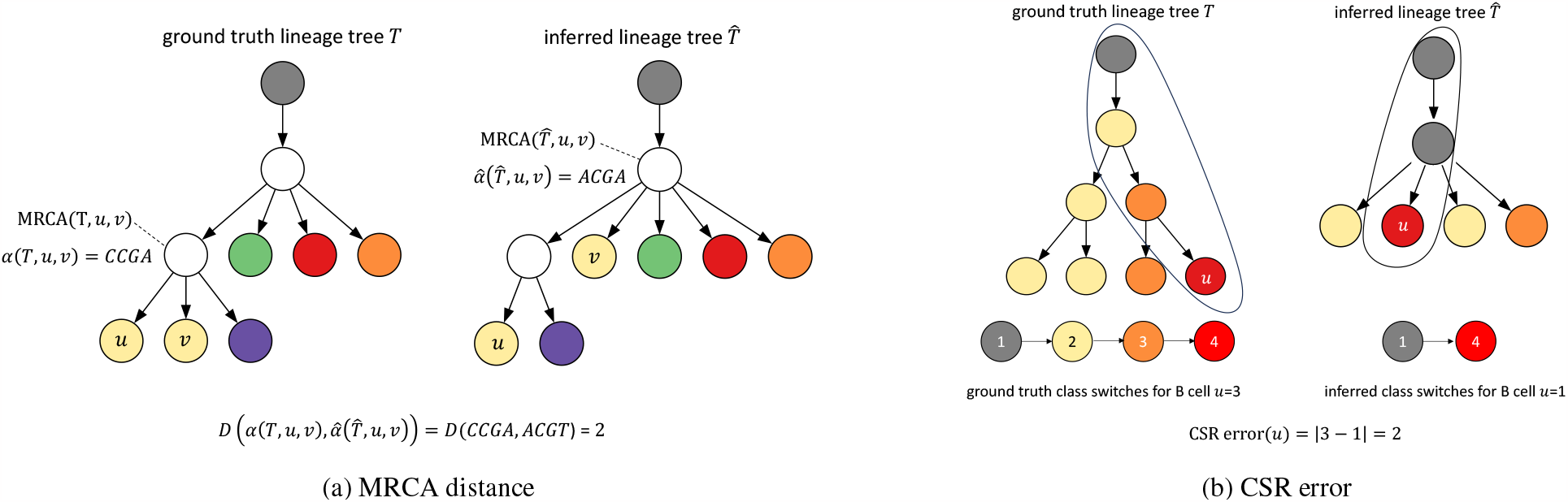
Performance metrics for B cell lineage tree inference. (a) An example calculation for MRCA distance leaves *u* and *v*. (b) An example calculation of CSR error for lineage *u*.

#### Kullback-Leibler (KL) divergence

To evaluate accuracy of isotype transition probability inference, we used *Kull-back–Leibler (KL) divergence* [27] to compare the inferred transition probability distribution 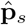 of each isotype *s* to the simulated ground truth distribution **p**_*s*_. KL divergence *D*_KL_ is defined as

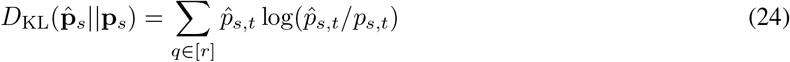

The lower the KL divergence, the more similar the two distributions.

### Normalized Robinson-Foulds (RF) distance

To assess the accuracy of topology of the inferred B cell lineage tree 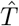 with respect to simulated ground truth tree *T*, we used *normalized Robinson-Foulds (RF) distance*. For this metric, we treat both trees as unrooted. For an unrooted tree, if you remove an edge (but not its endpoints), it defines a bipartition of the leaf set [43]. Doing this for every edge in tree *T* yields a set *B*(*T*) of bipartitions. RF distance is defined as the size of the symmetric difference between bipartitions *B*(*T*) and 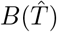 [28]. We then normalize this by the total number of bipartitions in each tree. Thus, normalized RF is computed as follows

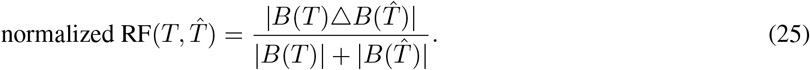

### Most Recent Common Ancestor (MRCA) distance

To assess the accuracy of the inferred ancestral sequence reconstruction 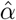 with respect to simulated ground truth *α*, we used a metric called *Most Recent Common Ancestor (MRCA) distanc*e introduced by Davidsen and Matsen [13]. For any two simulated B cells (leaves), the MRCA distance is the Hamming distance between the MRCA sequences of these two B cells in both the ground truth and inferred lineage trees. This distance is then averaged over all pairs of simulated B cells. A graphical depiction of this metric is show in Fig. S4a.

More formally,

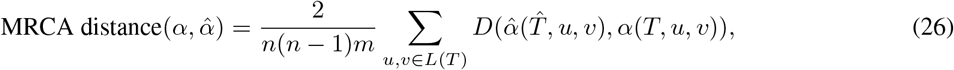

where in a slight abuse of notation *α*(*T, u, v*) is the sequence of the most recent common ancestor (MRCA) of nodes *u* and *v* in lineage tree *T* and *m* is the length of MSA.

### Class switch recombination (CSR) error

We assessed the accuracy of isotype inference by a new metric called *CSR error*, which is computed for each B cell *i* and clonotype *j* and is the absolute difference between the number of ground-truth class switches and inferred number of class switches that occurred along its evolutionary path from the root (Fig. S4b). Since dnaml, dnapars and IgPhyML do not infer isotypes for internal nodes, we pair these methods with the Sankoff algorithm [29] using *w*_*s,t*_ equals 1 if *s* = *t*, 0 if *s* < *t* and *∞* otherwise.

## E Supplementary Results

**Figure S5:**
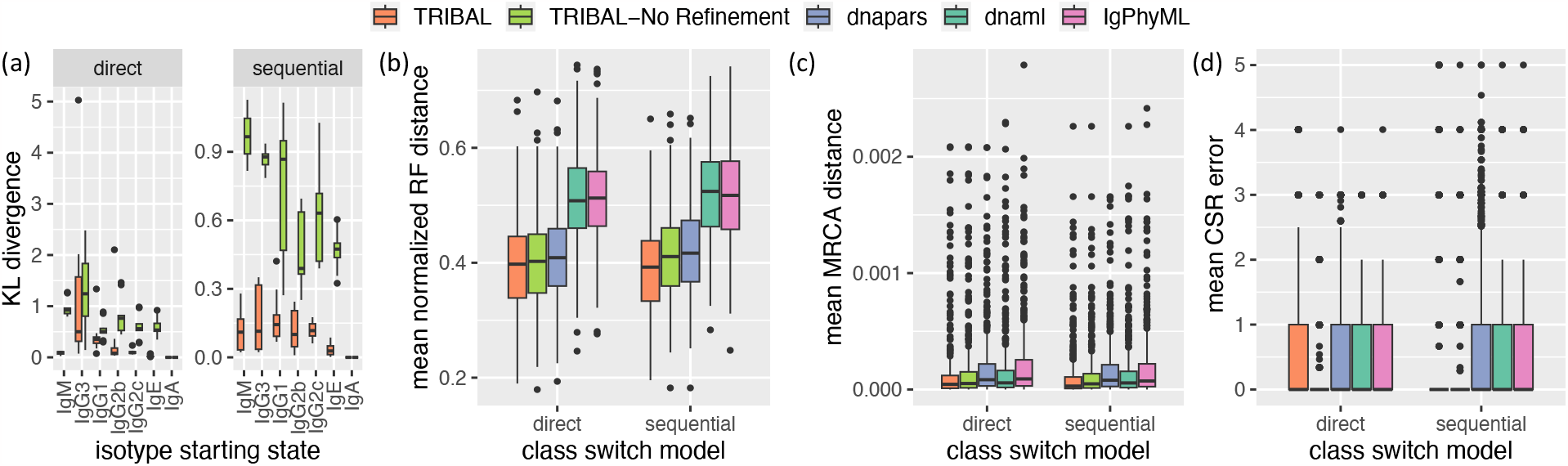
Simulations results for *k* = 75 clonotypes and *n* = 65 cells per clonotype.

**Figure S6:**
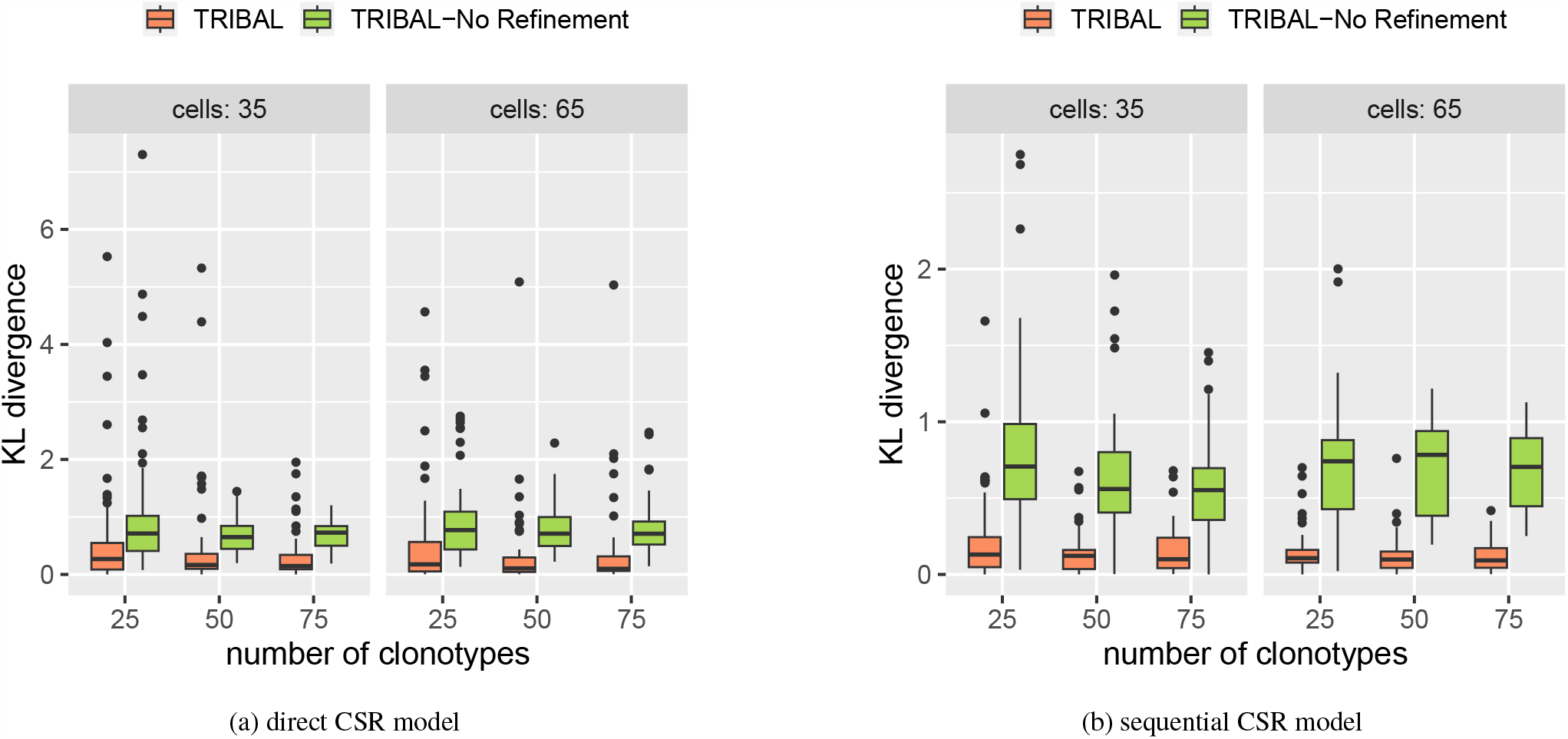
KL divergence from ground truth isotype transition probabilities aggregated over all isotype starting states, except IgA, by isotype starting state with varying the number *k* clonotypes, the number *n* of cells and CSR model.

**Figure S7:**
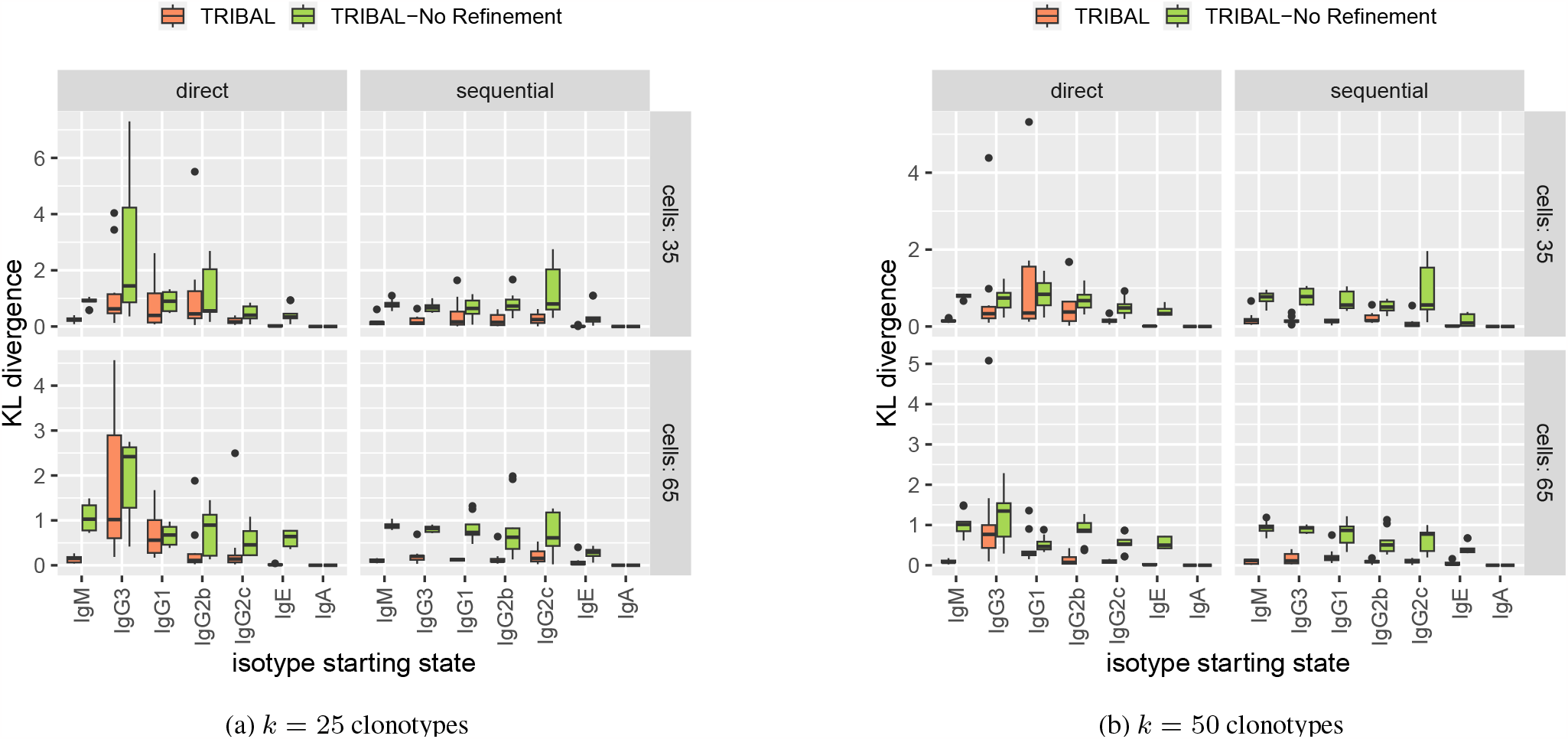
KL divergence from ground truth isotype transition probabilities by isotype starting state for *k∈ {*25, 50} clonotypes.

**Table S1:** Summary of NP-KLH mouse scRNA-seq datasets.

| dataset | clonotypes $k$ | total cells $n$ | median cells per clonotype | max cells per clonotype | median distinct isotypes per clonotype |
| --- | --- | --- | --- | --- | --- |
| NP-KLH-1 | 167 | 1776 | 7 | 89 | 3 |
| NP-KLH-2a | 70 | 537 | 6 | 32 | 2 |
| NP-KLH-2b | 58 | 357 | 5 | 21 | 2 |

### E.1 Average clade entropy for a leaf labeling

We describe a metric used to assess the average entropy contained within a leaf-labeling of the clades of a tree. First, we introduce some notation. Let Σ be an alphabet. Let clade *u* of tree *T* be the subtree *T*_*u*_ rooted at node *u*. Let *δ*(*u*) ⊆*L*(*T*) be the subset of leaves that are descendants of node *u*. Let *ℓ* : *L*(*T*)→Σ be a leaf labeling. Given a clade *u* and leaf-labeling *ℓ*, the entropy of a clade with respect to its leaf labels is defined as

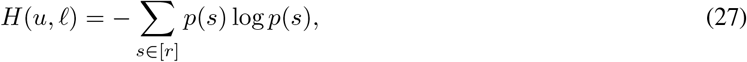

where *p*(*s*) =∑_*v∈δ*(*u*)_ **1**(*ℓ*(*v*) = *s*)*/*|*δ*(*u*)|. The average clade entropy 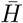 is computed over all clades except the leaves *L*(*T*) and the root *r* as follows

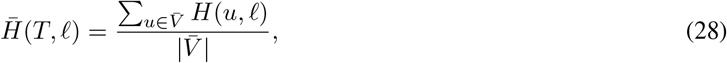

where 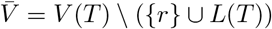 is the set of non-trivial clades.

**Figure S8:**
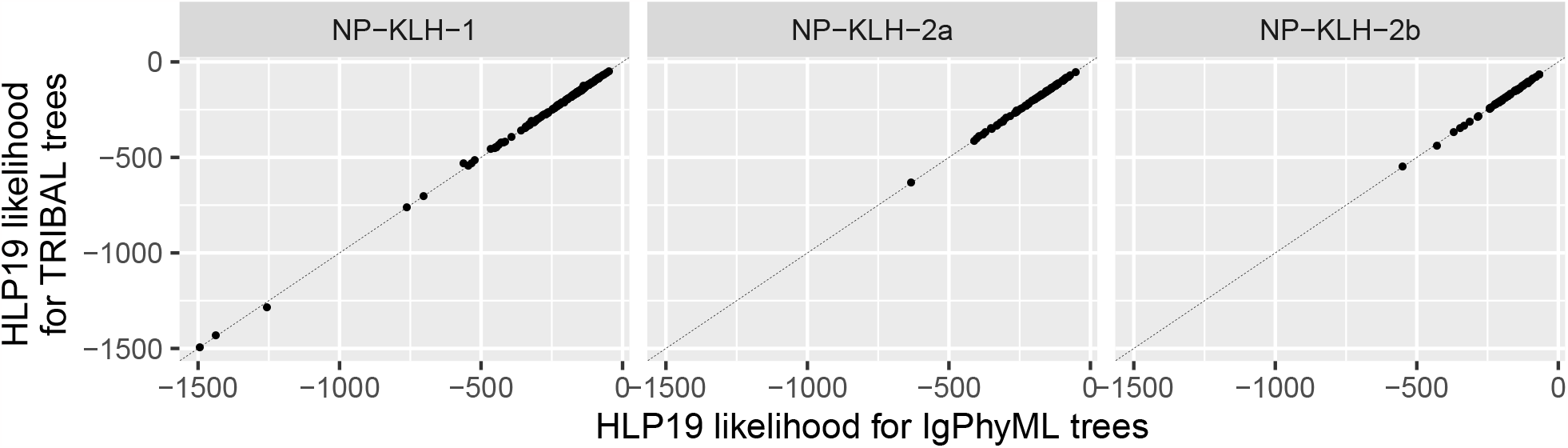
Comparison of HLP19 likelihood computed for IgPhyML and TRIBAL inferred B cell lineage trees for NP-KLH datasets.

**Figure S9:**
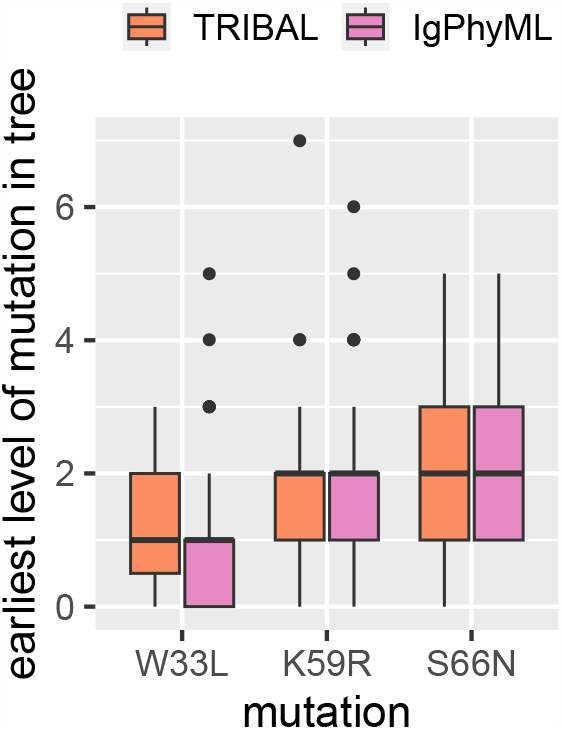
Earliest observed level of mutation in a B cell lineage tree. Level 0 represents the MRCA.

**Figure S10:**
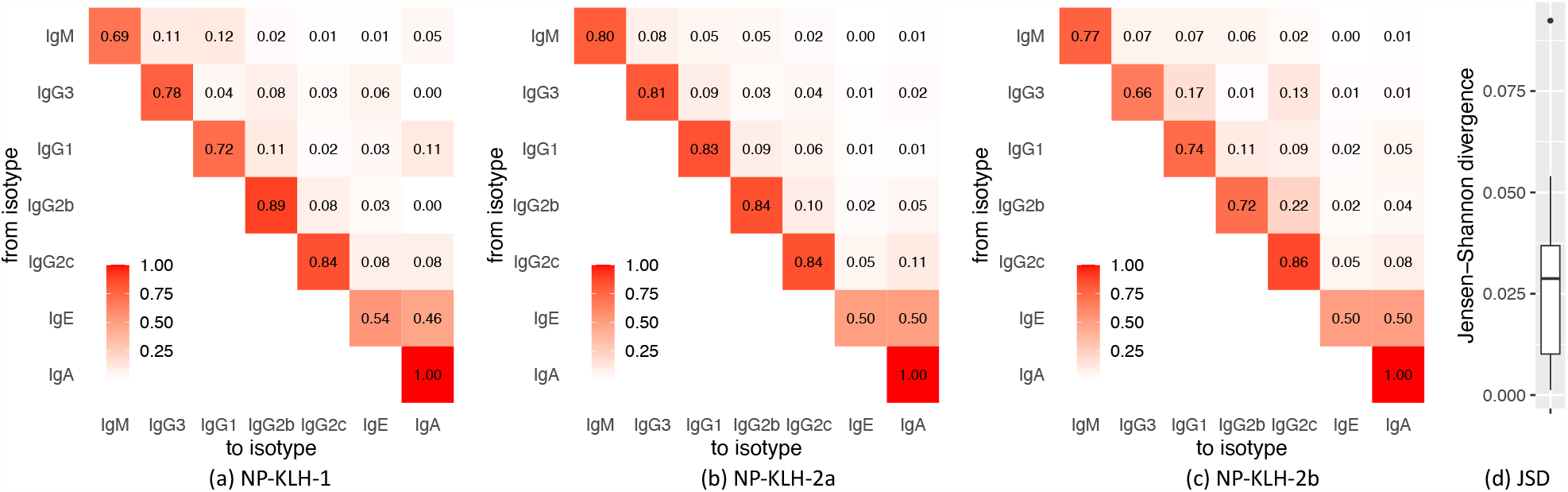
TRIBAL inferred isotype transition probabilities for NP-KLH. (a) Isotype transition probabilities for NP-KLH-1. (b) Isotype transition probabilities for NP-KLH-2a. (c) Isotype transition probabilities for NP-KLH-2b. (d) The distribution of Jensen-Shannon divergence (JSD) for pairwise comparisons of rows of the inferred isotype transition probabilty matrices for IgM through Ig2c. IgE was excluded from comparison due to a lack of observed B cells within each dataset to yield informative estimates. IgA is excluded as the inference of this row is trivial.

**Figure S11:**
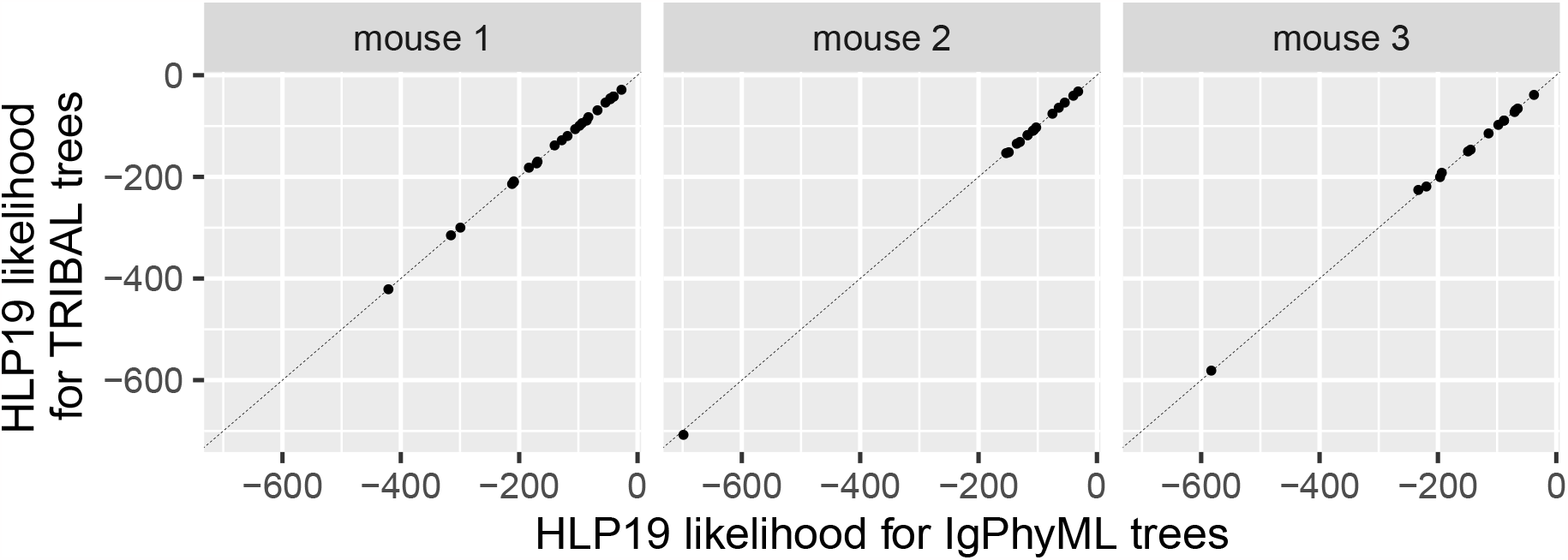
Scatterplot comparing HLP19 likelihood for IgPhyML trees to the HLP19 likelihood computed for TRIBAL trees for ABC datasets.

**Table S2:** Summary of ABC mouse scRNA-seq datasets.

| dataset | clonotypes $k$ | total cells $n$ | median cells per clonotype | max cells per clonotype | median distinct isotypes per clonotype |
| --- | --- | --- | --- | --- | --- |
| mouse 1 | 24 | 224 | 7.5 | 31 | 2 |
| mouse 2 | 15 | 218 | 7 | 81 | 3 |
| mouse 3 | 15 | 157 | 7 | 39 | 3 |

**Figure S12:**
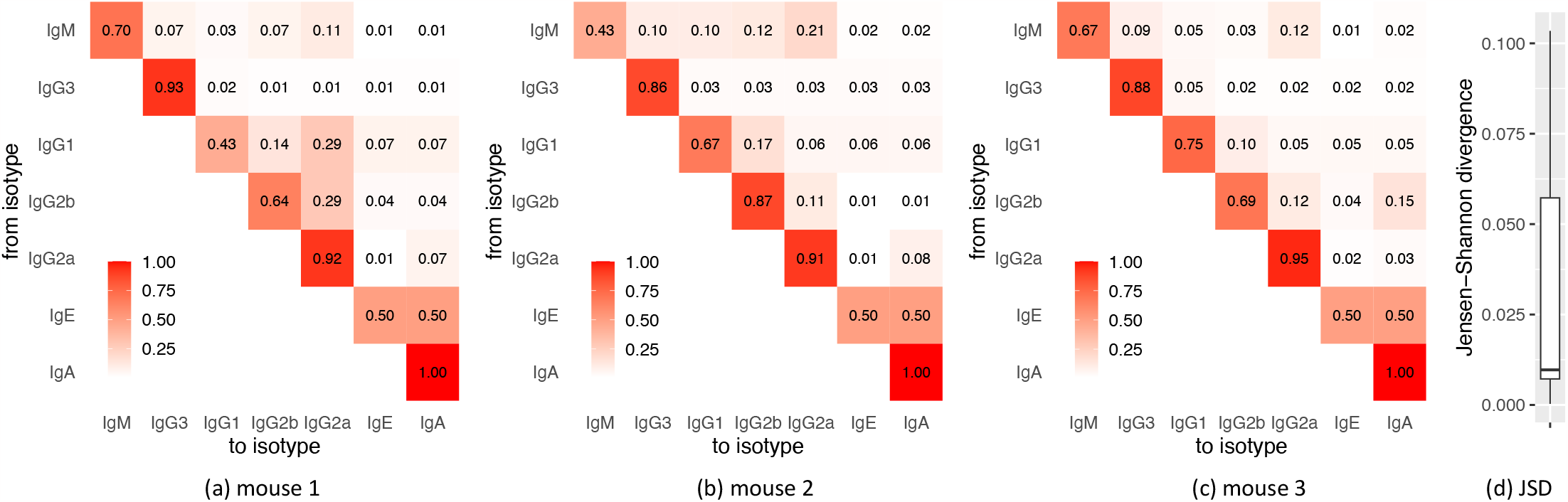
TRIBAL inferred isotype transition probabilities for ABC datasets. (a) Isotype transition probabilities for Mouse 1. (b) Isotype transition probabilities for Mouse 2. (c) Isotype transition probabilities for NP-Mouse 3. (d) The distribution of Jensen-Shannon divergence (JSD) for pairwise comparisons of rows of the inferred isotype transition probability matrices for IgM through Ig2c. IgE was excluded from comparison due to a lack of observed B cells within each dataset to yield informative estimates. IgA is excluded as the inference of this row is trivial.

## Footnotes

1 krdav/bcr-phylo-benchmark

## Notes

https://github.com/elkebir-group/TRIBAL

### Competing Interest Statement

The authors have declared no competing interest.

